# Blood-based detection of MMP11 as a marker of prostate cancer progression regulated by the ALDH1A1-TGF-β1 signaling mechanism

**DOI:** 10.1101/2024.07.16.603771

**Authors:** Ielizaveta Gorodetska, Vasyl Lukiyanchuk, Marta Gawin, Myroslava Sliusar, Annett Linge, Fabian Lohaus, Tobias Hölscher, Kati Erdmann, Susanne Fuessel, Angelika Borkowetz, Mark Reardon, Ananya Choudhury, Yasmin Antonelli, Aldo Leal-Egaña, Ayse Sedef Köseer, Uğur Kahya, Jakob Püschel, Daria Klusa, Claudia Peitzsch, Romy Kronstein-Wiedemann, Torsten Tonn, Christian Thomas, Piotr Widłak, Monika Pietrowska, Mechthild Krause, Anna Dubrovska

## Abstract

**Background:** Prostate cancer (PCa) is the second most common type of tumor diagnosed in men and the fifth leading cause of cancer-related death in male patients. The response of metastatic disease to standard treatment is heterogeneous. As for now, there is no curative treatment option available for metastatic PCa, and the clinical tests capable of predicting metastatic dissemination and metastatic response to the therapies are lacking. Our recent study identifies aldehyde dehydrogenases ALDH1A1 and ALDH1A3 as critical regulators of PCa metastases. Still, the exact mechanisms mediating the role of these proteins in PCa metastatic dissemination remain not fully understood, and plasma-based biomarkers of these metastatic mechanisms are also not available.

**Methods:** Genetic silencing, gene overexpression, or treatment with different doses of the retinoic acid (RA) isomers, which are the products of ALDH catalytic activity, were used to modulate the interplay between retinoic acid receptors (RARs) and androgen receptor (AR). RNA sequencing (RNAseq), reporter assays, and chromatin immunoprecipitation (ChIP) analysis were employed to validate the role of RARs and AR in the regulation of the transforming growth factor-beta 1 (TGFB1) expression. Gene expression levels of ALDH1A1, ALDH1A3, and the matrix metalloproteinase 11 (MMP11) and their correlation with pathological parameters and clinical outcomes were analyzed by mining several publicly available patient datasets as well as our multi-center transcriptomic dataset from patients with high-risk and locally advanced PCa. The levels of MMP11 protein were analyzed by enzyme-linked immunosorbent assay (ELISA) in independent cohorts of plasma samples from patients with localized or metastatic PCa and healthy donors, while plasma proteome profiles were obtained for selected subsets of PCa patients.

**Results:** We could show that ALDH1A1 and ALDH1A3 genes differently regulate TGFB1 expression in a RAR-and AR-dependent manner. We further observed that the TGF-β1 pathway contributes to the regulation of the MMPs, including MMP11. We have confirmed the relevance of MMP11 as a promising clinical marker for PCa using several independent gene expression datasets. Further, we have validated plasma MMP11 levels as a prognostic biomarker in patients with metastatic PCa. Finally, we proposed a hypothetical ALDH1A1/MMP11-related plasma proteome-based prognostic signature.

**Conclusions:** TGFB1/MMP11 signaling contributes to the ALDH1A1-driven PCa metastases. MMP11 is a promising blood-based biomarker of PCa progression.

## Background

Prostate cancer (PCa) is the second most common malignancy in men, accounting for about 14% of cancer diagnoses (1). If tumor growth is limited to the prostate, curative treatments are available, and the relative 5-year survival is close to 100% (2). Patients with localized PCa can be treated with curative intent by surgery or radiotherapy with or without androgen deprivation therapy (ADT). ADT combined with androgen receptor-targeted agents or taxane-based chemotherapy is given as a standard treatment to patients with metastatic PCa; however, after 18-24 months of treatment patients usually show disease progression toward castration-resistant prostate cancer (CRPC) (3). Patients with advanced PCa pose a high risk of bone metastases, which affect about 70%-90% of patients in the late stages of the disease, being a major cause of PCa morbidity and mortality (4, 5). When tumors spread beyond the prostate, long-term cure is often not possible anymore. The response of metastatic PCa to standard PCa treatment, such as ADT and radiotherapy, is heterogeneous. Tumor spread to bone is associated with lowering the relative 5-year survival to about 33% (2). As for now, there is no curative treatment option possible for PCa patients with metastatic disease, and clinically approved tests capable of predicting metastatic dissemination and metastatic response to the therapies are not available (6).

A high bone metastatic tropism of PCa cells is mediated by cues from the bone microenvironment, providing a favorable niche for cancer cell colonization, survival, and growth (7). The exact molecular mechanisms mediating metastatic spread and therapy resistance are poorly understood. Nevertheless, several signaling mechanisms have been implicated in bone metastatic dissemination. Transforming growth factor-beta 1 (TGF-β1) was reported to play a critical role in PCa progression by promoting epithelial-mesenchymal transition (EMT). TGF-β1 signaling orchestrates tumor invasion and angiogenesis by regulation of the matrix metalloproteinase (MMP) expression. MMPs are a large family of zinc-dependent endopeptidases mediating a proteolytic degradation of the extracellular matrix (ECM) and playing a pivotal role in cancer progression and metastatic spread (8). In PCa, TGF-β1 induces expression and secretion of MMP2 and MMP9 (9, 10). Both MMPs are regulated by androgen and serve as promising biomarkers and therapeutic targets in PCa (11). Once tumor cells are disseminated to bones, TGF-β1 plays a role in bone metastasis formation by modulating the interactions between PCa cells, osteoblasts, and osteoclasts (12, 13). Furthermore, TGF-β1 is an inducer of cancer cell reprogramming (14) and cancer stem cell (CSC) phenotype (15). In different types of tumors, TGF-β1-driven canonical Smad pathway and non-canonical TGF-β1 signaling have been shown to control the expression of the aldehyde dehydrogenase (ALDH) genes ALDH1A1 and ALDH2 and regulate a population of CSCs with high ALDH activity in a context-specific manner (16–18).

ALDH proteins are an enzyme superfamily involved in the oxidation of aldehydes to carboxylic acids, playing a crucial role in cellular detoxification, metabolism, and regulation of CSCs (19). The products of the enzymatic activity of ALDH proteins are retinoic acid (RA) isomers serving as ligands for retinoic acid receptors (RARs) and retinoid X receptors (RXRs). RARs and RXRs nuclear receptors bind to the RA response element (RARE) in the promoters of retinoid-responsive genes and trigger transcriptional activation in an RA-dependent manner (19–22). The all-trans RA (ATRA) is a clinically approved therapeutic agent for acute promyelocytic leukemia (APL). It has also been shown to have preclinical efficacy for other types of tumors (23). Previous studies on other types of tumors, including mesothelioma as well as breast and pancreatic cancer, revealed that ATRA and RARA act at multiple levels of the TGF-β1 signaling and regulate expression of the receptors, ligands, and signal transducers suggesting crosstalk of the TGF-β1 and retinoid signaling (24–27).

The ALDH1A1 and ALDH1A3 proteins have been identified as the primary isoforms accountable for ALDH activity in PCa cells and are involved in the production of RA from retinol (28). Our previous study revealed an opposite role of ALDH1A1 and ALDH1A3 genes in regulating PCa bone metastases. The distinct functions of these genes were attributed to their interplay with RAR-and androgen receptor (AR)-mediated expression of metastatic regulators such as polo-like kinase 3 (PLK3) (29).

Our current study further investigates the molecular mechanisms regulated by ALDH proteins that contribute to maintaining PCa metastasis-initiating cells. By genetic silencing of ALDH1A1 and ALDH1A3, RNAseq profiling, and chromatin immunoprecipitation (ChIP) analyses, we observed that ALDH1A1 and ALDH1A3 genes differently regulate TGFB1 expression in a RAR-and AR-dependent manner. We further demonstrated that the TGF-β1 pathway contributes to the regulation of the MMPs, including MMP11 and MMP26. We have confirmed the relevance of MMP11 as a potential clinical marker for PCa using several independent publicly available gene expression datasets and our multicenter gene expression analysis. Next, we have validated plasma MMP11 levels as a promising prognostic biomarker in patients with oligometastatic PCa. Finally, we employed proteomic profiling of plasma samples to reveal an ALDH1A1/MMP11-related prognostic signature.

## Methods

### Cell lines and culture condition

The human PCa cell lines PC3 (derived from bone metastasis), LNCaP (derived from lymph node metastasis), LNCaP-C4-2B (further named C4-2B, a bone metastatic derivative subline of the LNCaP cell line) and human embryonic kidney HEK293T cells were purchased from the American Type Culture Collection (Manassas, VA, USA) and cultured according to the manufacturer’s recommendations in a 37°C incubator in a humidified atmosphere with 5% CO_2_. The PC3 cell line was cultivated in DMEM medium; LNCaP and C4-2B cells in RPMI1640 medium. L-glutamine-free cell medium (PAN Biotech, Germany) was supplemented with 10% FBS (Capricorn Scientific, Germany) and 1 mM L-glutamine (Sigma-Aldrich, USA). HEK293T cells were used for luciferase reporter assay and cultured in DMEM medium supplemented with 10% FBS and 1 mM L-glutamine. All cell lines were subject to regular testing for mycoplasma contamination and genotyping using microsatellite polymorphism analyses.

### Preparation of 3D spheroids and tumor-like microcapsules

PCa spheroids were obtained by seeding 2-4x10^4^ cells in 24-well plates pre-coated with 250 μl of agarose (1.0% w/v). Tumor-like microcapsules were made using a Pocket-Microencapsulator device (patent pending, application number EP23207537.4). Microcapsules were made of 1.0% w/v alginic acid (Sigma-Aldrich, USA) and 1.0% w/v gelatin type A (Sigma-Aldrich, USA). These polymers were dissolved in a buffer containing 0.1 M HEPES (pH 7.4) (Sigma-Aldrich, USA), 1.0% w/v NaCl (Carl Roth, Germany), and 1.0% v/v penicillin-streptomycin (Sigma-Aldrich, USA). The blend was then filtered using a 0.45 μm pore diameter filter and loaded with PCa cells with a density of 500,000 cells/ml. 0.6 M CaCl_2_ (Carl Roth, Germany) was used to crosslink the alginate-gelatin microcapsules, which were then dissolved in 0.1 M HEPES (pH 7.4) containing 1.0% w/v NaCl. On day 3, PCa cells were isolated from the microcapsules by immersing them for 1 min in a solution containing 22.5 mM of sodium citrate dihydrate (Carl Roth, Germany), 60 mM EDTA (Sigma-Aldrich, USA), and 150 mM NaCl (pH 7.4), following an already published protocol (30). After that, cells were collected by centrifugation (1000 rpm, 5 min).

### ATRA and TGF-β1 treatment

ATRA (Cayman Chemical, USA) was dissolved in DMSO (Fisher Scientific, USA), and corresponding concentrations of DMSO were used as controls. The cells were serum starved in DMEM or RPMI with 3% FBS for 24 h followed by treatment with 10, 25, or 50 μM of ATRA or 9CisRA (Cayman Chemical, USA) for 48 h. For TGFβ1 treatment experiments, cells were incubated with serum-reduced (5% FBS) growth medium with or without TGF-β1 (Miltenyi Biotec, Germany) at the concentration of 5 ng/ml. After 48 h in the incubator, cells were used for functional assays or RNA isolation by NucleoSpin RNA isolation kit (Macherey-Nagel, Germany). In total, at least three independent biological repeats were performed with cells at different passages.

### siRNA-mediated gene silencing

The cells were grown until confluency of 60-80% in complete medium and transfected by specific siRNA using Xfect RNA transfection reagent (Takara Bio, USA) according to the manufacturer’s instructions. Cells transfected with unspecific siRNA (scrambled siRNA or siSCR) were used as a negative control in all knockdown experiments. Cells were harvested 48 h after transfection. The RNA duplexes were synthesized by Eurofins and used as a pool of 2-4 duplexes per target. The siRNA sequences used in the study are described in Table S1.

### RNA isolation, cDNA synthesis, and qPCR

RNA from PCa cells was isolated by NucleoSpin RNA isolation kit (Macherey-Nagel, Germany). Reverse transcription was done using the PrimeScript™ RT reagent Kit (Takara Bio, USA) according to the manufacturer’s recommendation. Quantitative real-time polymerase chain reaction (qPCR) was carried out using the TB GreenTM Premix Ex TaqTM II (Takara Bio, USA) according to the manufacturer’s protocol for a total reaction volume of 20 μl. The qPCR cycling program was set on a StepOnePlus system (Applied Biosystems, USA): 95°C for 10 min, 40 cycles: 95°C for 15 s, 60°C for 60 s, 72°C for 60 s followed by a melt curve to 95°C in steps of 0.3°C. All experiments were conducted using at least three technical replicates. The expression of ACTB and RPLP0 mRNA was used for data normalization. The primers used in the study are listed in Table S1.

### Oris migration assay

Oris migration assay (Platypus Technologies, USA) was used to validate the ability of the cells to migrate *in vitro.* Firstly, 200,000 cells/well were seeded in 6-well plates and were incubated for 24 h at 37°C and 5% CO_2_. After 24 h, cells were trypsinized and plated in 96-well collagen I-coated plates at a density of 30,000 cells/well. Silicon stoppers (Platypus Technologies, USA) were inserted into the wells to keep the middle of the well free of cells. After 24 h, stoppers were removed, and cells were scanned using the Celigo S Imaging Cell Cytometer (Nexcelom) to define the pre-migration area (t = 0 h). The plate was then incubated at 37°C and 5% CO_2_ to permit the migration and detection of the invaded middle zone of the wells. After 24 h, the plate was scanned again. The pictures were analysed by ImageJ software and the area invaded by cells within this time was compared.

### Chromatin immunoprecipitation (ChIP)

2x10^6^ LNCaP cells were plated in 150 mm Petri dishes and were treated with 50 µM of ATRA the next day. 48 h after the treatment started, a crosslinking was performed by incubation with 1% formaldehyde for 10 minutes at 37°C. Cells were collected in PBS containing a protease inhibitor cocktail (#5872, Cell Signaling Technology, USA). DNA fragmentation step was conducted using Micrococcal Nuclease (#10011, Cell Signaling Technology, USA), and nuclei were transferred to SDS lysis buffer. ChIP experiments were performed with the ChIP Assay Kit (#17-295, Merck Millipore, Germany) according to the manufacturers instructions. The samples were incubated with primary antibodies against AR (#5153) and RARA (#62294) or control Rabbit IgG antibody (#3900) (all Cell Signaling Technology, USA) overnight at 4°C. On the next day, DNA-protein-antibody complexes were precipitated using Agarose beads, and crosslinks were reversed at 65°C for 4 h. The DNA fragments were purified using the QIAquick PCR purification kit (Qiagen, Germany). For qPCR detection of immunoprecipitated DNA fragments, primers were designed to cover different promoter regions of the presumable AR or RARA target genes, which contained putative AR or RARA binding sites (predicted by Eukaryotic Promoter Database, https://epd.epfl.ch//index.php). The significance of DNA fragment yield after AR or RARA antibody precipitation compared to control IgG was calculated using a paired t-test.

### Plasmid DNA and cell transfection

The DNA plasmids used in this study include pcDNA3.1 (Invitrogen, USA), pEGFP-N3 (Clontech, USA), pcDNA3.1-hRARα (31) (#35397, Addgene, USA), pGL3-TGFb1(32) (#101762, Addgene, USA). Plasmids were amplified in E. coli strain ΔH5α (Invitrogen, USA) and isolated using NucleoBond Xtra Midi (Macherey-Nagel, Germany) according to the protocol from the manufacturer. For the gene reporter assay, HEK293T cells were transfected using the calcium phosphate method. Transfection with 1.25 μg pEGFP-N3 plasmid was used as a control of transfection efficacy. Cells were treated with various concentrations of ATRA (10 μM, 25 μM, and 50 μM) and DMSO as a control. Luciferase activity (Promega, USA) was measured 48 h after the treatment started. For the RARA overexpression, LNCaP cells were transfected using Xfect transfection reagent (Takara Bio, USA) according to the manufacturer’s recommendations. Cells were treated with 50 μM of ATRA and DMSO as a control. Cells transfected with an empty plasmid DNA were used as a negative control. 48 h after the treatment started, cells were collected for RNA isolation.

### Fluorescence and luminescence measurements

GFP fluorescence was measured by fluorescence reader SpectraMax iD3 (Molecular Devices, USA) at 525 nm following 485 nm excitation. Luciferase activity was measured by Bright-GloTM Luciferase Assay System (Promega, USA). Luminescence was measured by luminescence reader SpectraMax iD3 after 2 min of shaking to allow complete cell lysis.

### Clinical specimens

Clinical specimens were collected with informed consent from all subjects. The ethical approvals for these retrospective analyses were granted by the local Ethics Committees and Institutional Review Boards of the Faculty of Medicine, Technische Universität Dresden, Germany (EK194052014 and EK3012017) and the University of Manchester, Manchester Cancer Research Centre (#15/NW/0559).

Blood samples (n = 31) were collected from patients with androgen-sensitive oligometastatic PCa treated with metastasis-directed local ablative external beam radiation therapy (EBRT) (conventionally fractionated 25 x 2 Gy or hypofractionated 3 x 10 Gy) at the Department of Radiotherapy and Radiation Oncology at the University Hospital Carl Gustav Carus, Dresden (ClinicalTrials.gov Identifier: NCT02264379) between 2014 and 2021 and from healthy donors of above 55 years old (n = 5) collected at the Institute for Transfusion Medicine Dresden and serving as control. The characteristics of patients are provided in Table 1. Additional marker validation was conducted using plasma samples (n = 43) from patients with localized or metastatic PCa (hormone-sensitive PCa or CRPC) treated with radical prostatectomy (primary PCa) or taxane-based chemotherapy (metastatic PCa), respectively, at the Department of Urology of the University Hospital Carl Gustav Carus, Dresden. The characteristics of patients are provided in Table 2.

**Table 1.** Characteristics of patients with androgen-sensitive, oligometastatic prostate cancer (Oli-P study cohort) that received ablative radiotherapy with curative intend to metastatic lesions. Patients were treated at the Department of Radiotherapy and Radiation Oncology of the University Hospital Carl Gustav Carus, Dresden (ClinicalTrials.gov Identifier: NCT02264379) between 2014 and 2021.

| Parameter | Number of patients | % |
| --- | --- | --- |
| **ISUP grade** |  |  |
| High ($\geq 4$) | 10/31 | 32% |
| **NCCN risk group** |  |  |
| Low | 0/31 | 0% |
| Intermediate | 9/31 | 29% |
| High, very high, local advanced | 22/31 | 71% |
| **Metastases** |  |  |
| Lymph node | 21/31 | 68% |
| Bone | 6/31 | 19% |
| Both | 4/31 | 13% |
| **Radiotherapy to metastases** |  |  |
| Normo-fractionation (25 x 2 Gy) | 7/31 | 23% |
| Hypo-fractionation (3 x 10 Gy) | 24/31 | 77% |
| Parameter | Mean | Range |
| Age (years) | 73 | 53-84 |
| Gross tumor volume (GTV) (ccm) | 3.0 | 0.1-11.0 |
| PSA before radiotherapy (ng/ml) | 3.02 | 0.69-8,78 |

GTV - gross tumor volume; the sum of one to three metastatic lesions.
ISUP - International Society of Urological Pathology
NCCN - National Comprehensive Cancer Network

**Table 2.** Characteristics of patients with (A) localized PCa treated by radical prostatectomy (n = 14) or (B) with metastatic PCa (hormone sensitive PCa or CRPC; n = 29) treated with taxane-based chemotherapy (docetaxel or cabazitaxel). The patients were treated at the Department of Urology of the University Hospital Carl Gustav Carus, Dresden.

| Parameter | Number of patients | % |
| --- | --- | --- |
| **ISUP grade** |  |  |
| High ($\geq 4$) | 10/14 | 71 |
| **Metastases** | 6/14 | 43 |
| Lymph node | 6/14 | 43 |
| Bone | 0/14 | 0 |
| Both | 0/14 | 0 |
| **Treatment** | radical prostatectomy | 100 |

| Parameter | Mean | Range |
| --- | --- | --- |
| Age (years) | 65.2 | 51.8-74.6 |
| PSA before treatment (ng/ml) | 19.70 | 4.49-80.59 |

**B)**
| Parameter | Number of patients | % |
| --- | --- | --- |
| **ISUP grade** |  |  |
| High ($\geq 4$) | 23/29 | 79 |
| **Metastases** | 29/29 | 100 |
| Lymph node | 19/29 | 66 |
| Bone | 27/29 | 93 |
| Both | 17/29 | 59 |
| **Treatment** | Taxane-based chemotherapy | 100 |

| Parameter | Mean | Range |
| --- | --- | --- |
| Age (years) | 68.5 | 51.6-92.9 |
| PSA before chemotherapy (ng/ml) | 285.5<br>5 unknown | 0.2-3134.0 |

For the evaluation of gene expression in PCa biopsies, the study included formalin-fixed, paraffin-embedded (FFPE) specimens from patients with high-risk and locally advanced PCa (pre-treatment PSA > 20 ng/ml, combined Gleason score ≥ 7 or clinical T-stage ≥ T2), treated at two UK cancer centers between 2006 and 2017. Patients were treated initially with ADT, then concurrently with one of three radiotherapy regimens delivered to prostate: conventionally fractionated EBRT (74 Gy, 37 fractions), n = 127; EBRT (37 Gy, 15 fractions) with a 12.5-15 Gy high-dose-rate (HDR) brachytherapy boost, n = 214; or hypofractionated EBRT (60 Gy, 20 fractions), n = 91. The cohort characteristics are described in Table 3. The gene expression analysis was conducted with Clariom^TM^ S Assay HT human microarray (Thermo Fisher Scientific, Massachusetts, US).

**Table 3.** Summary table of baseline clinical characteristics and biomarker scores for the cohort used for whole-genome transcriptomic analysis.

|  | **EBRT - Hypo.;**<br>60 Gy, 20 fractions<br>(n = 91) | **EBRT - HDR**<br>**boost;**<br>37 Gy, 15<br>fractions with a<br>12.5-15 Gy HDR<br>brachytherapy<br>boost<br>(n = 214) | **EBRT - Conv.;**<br>74 Gy, 37<br>fractions)<br>(n = 127) |
| --- | --- | --- | --- |
| **Age (years) – median**<br>**(min, max)** | 68 (51, 82) | 66 (46, 79) | 71 (53, 79) |
| *Missing* | 1 | 1 | 1 |
| **PSA (ng/ml) –**<br>**median (min, max)** | 19.3 (1.9, 180.0) | 20.8 (3.6, 225.0) | 17.5 (1.8, 234.0) |
| *Missing* | 5 | 5 | 1 |
| **ISUP grade**<br>**(n)** |  |  |  |
| 1 | 2 | 5 | 6 |
| 2 | 28 | 39 | 40 |
| 3 | 22 | 68 | 30 |
| 4 | 13 | 36 | 14 |
| 5 | 26 | 66 | 36 |
| **T stage (n)** |  |  |  |
| 1 | 6 | 6 | 0 |
| 2 | 31 | 66 | 8 |
| 3 | 51 | 139 | 111 |
| 4 | 2 | 1 | 0 |
| *Missing* | 1 | 2 | 8 |
| **Tumor (%) – median**<br>**(min, max)** | 50 (5, 95) | 65 (5, 100) | 60 (5, 100) |
| **ADT duration**<br>**(months) – median**<br>**(min, max)** | 27 (2, 39) | 18 (3, 99) | 24 (3, 36) |
| *Missing* | 0 | 4 | 2 |
| **CAPRA risk score (n)** |  |  |  |
| Low (0 – 2) | 0 | 0 | 2 |
| Intermediate (3 – 5) | 17 | 25 | 22 |
| High (6 – 10) | 74 | 189 | 103 |

ADT – androgen deprivation therapy; CAPRA – Cancer of the Prostate Risk Assessment; Conv – conventional radiotherapy; EBRT – external beam radiotherapy; HDR – high dose rate; Hypo – hypofractionated radiotherapy; PSA – prostate-specific antigen

### Enzyme-linked immunosorbent assay (ELISA)

The blood plasma of PCa patients and healthy donors was harvested with Biocoll (Bio&Sell, Germany) density gradient centrifugation along with the isolation of circulating tumor cells and stored at -80°C. The concentration of soluble, human MMP11 was measured using the MMP11 ELISA Kit (Antibodies online, ABIN6574149) and TGF-β1 by TGF beta 1 ELISA Kit (Abcam, ab108912, United Kingdom) according to manufacturers’ instructions within pre-diluted plasma aliquots (1:2).

### Mice and in vivo tumorigenicity assays

In vitro HNSCC tumor propagation was performed as described earlier (33). In brief, Cal33 RR HNSCC cells were embedded in 100 μl of DMEM/Matrigel mixed as 1:1 and injected subcutaneously into the hind legs of 7 to 12-weeks-old NMRI (nu/nu) mice. The immunosuppression was achieved by total body irradiation with 4 Gy of 200 kV X-rays (0.5 mm Cu filter, 1 Gy/min) 1 day before injections. The blood was taken from tumor-bearing mice by a cardiac puncture and collected using EDTA-coated tubes. The blood samples were centrifugated at 1200 rpm, 4°C for 15 min to separate plasma. The animal facility and the experiments were performed according to the institutional guidelines and the German animal welfare regulations (protocol number TVV 51/2022). The control plasma from 5 healthy NMRI (nu/nu) mice was purchased from Charles River (Germany).

### Mass spectrometry-based profiling of plasma proteome

Ten µL of plasma was added to 100 µL of lysis buffer (0.1M Tris-HCl pH 8.0, 0.1M DTT, 4% SDS), vortexed for 5 s (2000 rpm) and spun down shortly then boiled at 99°C for 10 min and left to cool down on a bench. An aliquot containing 100 µg of total protein was then subjected to Filter-Aided Sample Preparation according to the previous study (34, 35). Briefly, proteolytic digestion was performed using Trypsin/Lys-C Mix (Promega) with the enzyme-to-protein ratio of 1:25 (m/m), 37 °C, 18h, then resulting peptides were fractionated into 6 fractions using an anion exchanger (SAX StageTips); an aliquot of each fraction was subjected to nLC-MS/MS measurements. Q Exactive Plus mass spectrometer coupled with RSLC UltiMate 3000 nano-liquid chromatograph and Nanospray Flex ion source (all from Thermo Scientific) were used for measurements. Peptides from each fraction were separated on a reverse-phase Acclaim PepMap RSLC nanoViper C18 column (75μm × 25cm, 2μm granulation) using 80% acetonitrile in 0.1% formic acid gradient (from 3 to 60% as follows: 3-8% B during 7 minutes; 8-35% B 70 minutes; 35-60% B 10 minutes) at 35°C and a flow rate of 300nL/min (total run time: 110min). The spectrometer was operated in data-dependent MS/MS mode with survey scans acquired at a resolution of 70,000 at m/z 200 in MS mode, and 17,500 at m/z 200 in MS2 mode. Spectra were recorded in the scanning range of 300– 2000 m/z in the positive ion mode. Higher energy collisional dissociation (HCD) ion fragmentation was performed with normalized collision energies set to 25. Protein identification was performed using a reviewed Swiss-Prot human database (released May 2022 containing 11385168 residues and 20330 sequence entries) with a precision tolerance 10 ppm for peptide masses and 0.02 Da for the fragment ion masses. All raw data obtained for each dataset were imported into Protein Discoverer version 2.3.0.522 (Thermo Fisher Scientific) <Thermo raw files> for protein identification and quantification (Sequest engines were used for database searches). Protein was considered as positively identified if at least two peptides per protein were found by both search engines and a peptide score reached the significance threshold FDR = 0.01 (assessed by the Percolator algorithm); a protein was further considered as “present” if detected in at least one sample of a given type. The abundance of identified proteins was estimated in Proteome Discoverer using the Precursor Ions Area detector node, which calculates the abundance of a given protein based on the average intensity of the three most intensive distinct peptides for this protein, with further normalization to the total ion current (TIC). The high resolution mass spectrometry-based proteomic data have been deposited to the ProteomeXchange Consortium via the PRIDE (https://www.ebi.ac.uk/pride) (36, 37) partner repository with the dataset identifier PXD052973.

### Analysis of the patient gene expression datasets

The publicly available TCGA PRAD (38), Metastatic PCa SU2C/PCF Dream Team (39), MSKCC (40), DKFZ (41), SMMU (41) and Broad/Cornell (42) datasets were accessed via cBioportal (https://www.cbioportal.org/). For Kaplan-Meier survival analysis, the biochemical recurrence-free survival (BRFS) time (TCGA and DKFZ cohorts), the disease-free survival (DFS) time (MSKCC cohort), or the overall survival (OS) time (SU2C cohort) or metastasis-free survival (MFS) time (Manchester dataset, Table 3) and the PSA increase above nadir (Oli-P cohort, Table 1) were used as clinical endpoints. For the TCGA cohort, BRFS was determined based on provided "Days to PSA" and "Days to biochemical recurrence first" data. The patient groups were defined by the optimal cutoff scan procedure. The best cutoff for survival analysis and log-rank test p-values was determined using the R2 platform (https://hgserver1.amc.nl/cgi-bin/r2/main.cgi). We did not use Bonferroni correction for multiple testing since it assumes independence between multiple tests, which is not applicable for the best cutoff scan procedure.

### Statistical analysis

The results of the cell migration, ChIP analysis, luciferase reporter assay, ELISA, and relative gene expression determined by qPCR were analyzed by a paired two-tailed t-test. A significant difference between the conditions was defined as *p < 0.05; **p < 0.01; ***p < 0.001. The correlation of gene expression levels was calculated using the Pearson or Spearman (for nonparametric data) correlation coefficient. A statistical analysis for comparing the MMP11 expression levels in normal, primary tumor and metastatic tissues was performed by one-way ANOVA followed by posthoc Tukey HSD test. The diagnostic value of plasma MMP11 levels was evaluated by receiver operating characteristic (ROC) curve analysis. The diagnostic parameters, including sensitivity, specificity, accuracy, cut-off value, and area under the ROC curve (AUC) with 95% confidence interval (CI) and p-value, were determined by the ROC Plotter online tool (43) (https://rocplot.org/). An easyROC web-tool (44) was used for the simultaneous estimation of the prognostic potential for 20 MMP genes described in Figure 4 A (http://biosoft.erciyes.edu.tr/app/easyROC/).

## Results

### ALDH1A1 and ALDH1A3 differentially regulate cell migration depending on the AR status

Metastasis formation is a multistep process, and the acquisition of a migratory phenotype by tumor cells is one of the first steps of the metastatic cascade (45). The cell exclusion zone (Oris) migration assay was used to explore the role of ALDH genes in cell migration. For this analysis, we performed siRNA-mediated knockdown of ALDH1A1 and ALDH1A3 by using a siRNA pool and cells transfected with scrambled (Scr) siRNA as control. This experiment showed that ALDH1A1 and ALDH1A3 knockdown differently affect the migration properties of the PCa cells depending on their androgen sensitivity. While ALDH1A1 downregulation decreased migration in androgen-dependent LNCaP cells, it did not significantly affect migration in the LNCaP-derived C4-2B cells, which are AR-positive but androgen-independent, and even had a significant pro-migratory effect on the AR-negative PC3 cell line. Of note, ALDH1A3 knockdown decreased the migration rate only in PC3 cells (Figure 1A, Figure S1A).

**Figure 1.**
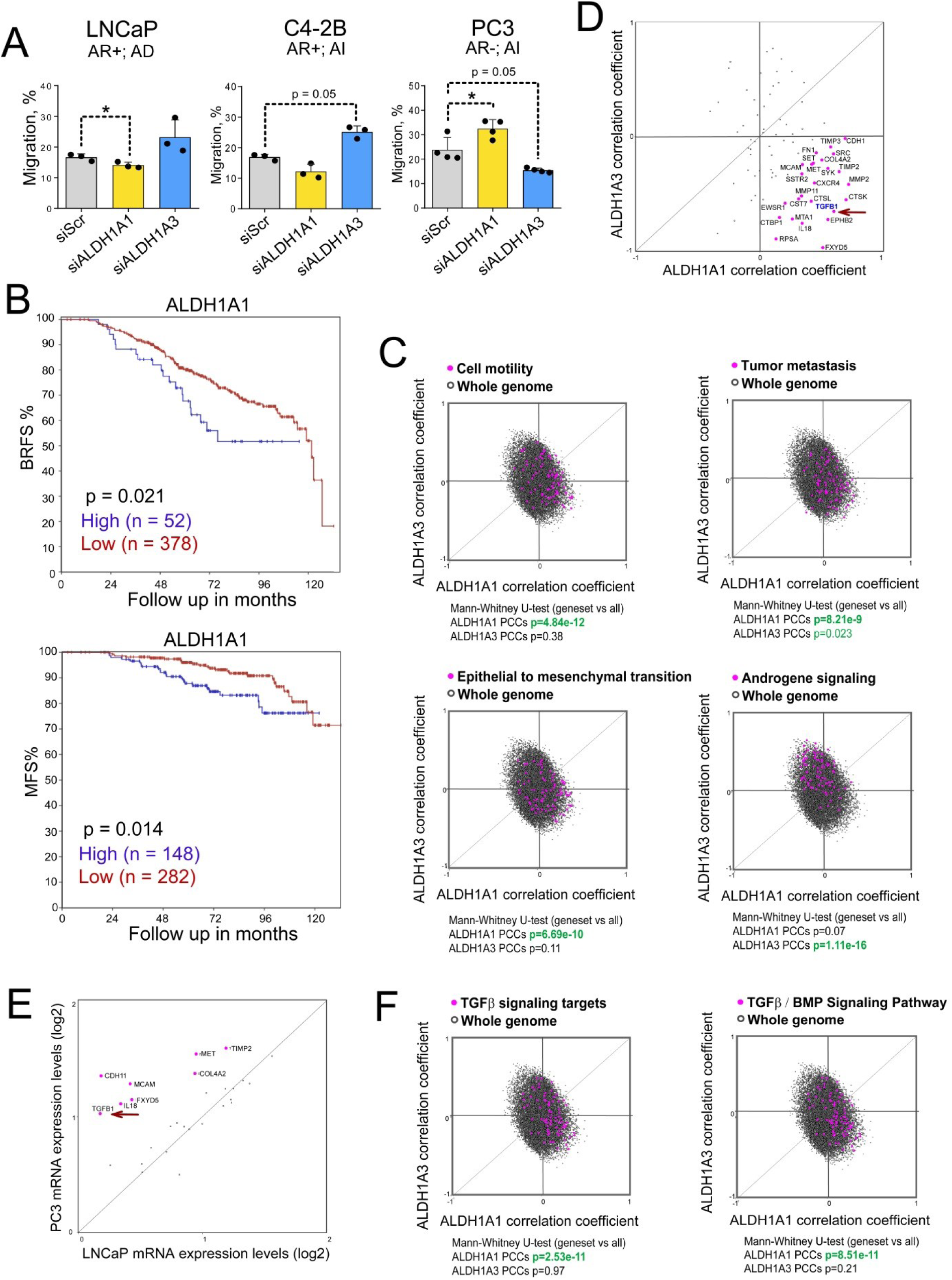
ALDH1A1 and ALDH1A3 differentially regulate migration depending on the AR status. (**A**) Analysis of the relative cell migration of LNCaP, C4-2B, and PC3 cells upon knockdown of ALDH1A1 or ALDH1A3 expression using Oris migration assay. Cells transfected with scrambled siRNA (siScr) were used as a control. Cell migration was analyzed 24 h after cell plating. AR+: androgen receptor positive; AR-: androgen receptor negative; AD: androgen dependent; AI: androgen independent; n = 3; Error bars = SD; *p < 0.05. (**B**) The Kaplan-Meier analyses of the biochemical recurrence-free survival (BRFS) and metastasis-free survival (MFS) for patients with high-risk and locally advanced PCa treated with ADT and then concurrently with one of three radiotherapy regimens as described in Table 3. The patients were stratified by the most significant cut-off for ALDH1A1 expression levels. (**C**) Correlation of mRNA expression levels of ALDH1A1 and ALDH1A3 genes and gene sets related to cell motility, tumor metastasis, EMT and androgen signaling in the TCGA PRAD patient cohort (n = 490). The gene lists are provided in Table S2. (**D**) Correlation of mRNA expression levels of ALDH1A1 and ALDH1A3 genes and metastasis-related genes in the TCGA PRAD patient cohort (n = 490). The arrow indicates a correlation with TGFB1. (**E**) Levels of mRNA expression of the metastasis-related genes which highly correlate with ALDH1A1 expression (as in panel D) in PC3 and LNCaP cell lines. The arrow indicates the level of TGFB1 expression. (**F**) Correlation of mRNA expression levels of ALDH1A1 and ALDH1A3 genes and gene sets related to the TGFβ1 signaling targets and TGFβ1 / BMP signaling targets in the TCGA PRAD patient cohort (n = 490). The gene lists are provided in Table S2.

Next, we analyzed a possible correlation of ALDH1A1 and ALDH1A3 gene expression with BRFS and MFS in a retrospective multicenter cohort including 432 patients diagnosed with locally advanced PCa (Manchester dataset). All patients were treated initially with ADT, then concurrently with one of three radiotherapy regimens delivered to the prostate, as described in Table 3. This analysis confirmed that high ALDH1A1 expression is associated with lower BRFS and MFS, whereas no significant association was found for ALDH1A3 expression (Figure 1B, Figure S1B). These observations are consistent with our previous findings that ALDH genes differently contribute to the clinical outcomes and metastatic dissemination (29). To further explore the role of ALDH genes in migration and metastasis, we used the TCGA gene expression dataset to analyze the potential correlation of ALDH1A1 and ALDH1A3 with gene signatures attributed to different molecular pathways. This analysis revealed a significant positive correlation of ALDH1A1 with several gene sets related to cancer progression and spread, including genes related to tumor cell motility, metastasis, EMT, ECM and adhesion molecules, angiogenesis, and osteogenesis. On the other hand, ALDH1A3 was strongly associated with the expression of the AR signaling targets consistent with our and other previous observations that ALDH1A3 is an AR transcriptional target (29) (Figure 1C, Figure S1C, Table S2).

Next, we checked if ALDH1A1 and ALDH1A3 expression correlates with 84 metastasis-related genes in the PCa TCGA cohort (n = 490). This analysis identified 25 genes having the highest positive correlation with ALDH1A1 and a negative correlation with ALDH1A3, with the TGFB1 gene being one of the top candidates (Figure 1D). Analysis of the expression levels of these metastasis-related genes in PC3 and LNCaP PCa cells identified TGFB1 as one of the genes highly expressed in bone metastasis-derived PC3 cells, but not in lymph node metastasis-derived LNCaP cells (Figure 1E). Indeed, genes attributed to the TGFβ / BMP signaling pathway highly correlated with ALDH1A1 but not with ALDH1A3 (Figure 1F). These analyses suggest a potential role of the interplay between ALDH1A1 and TGFB1 in metastatic development.

### ALDH1A1 and ALDH1A3 regulate TGFB1 gene expression through RA-dependent mechanism

We then analysed whether siRNA-mediated depletion of ALDH1A1 and ALDH1A3 affects TGFB1 gene expression. We found that ALDH1A1 downregulation decreased TGFB1 gene expression in LNCaP and C4-2B cells, however it did not regulate TGFB1 expression in PC3 cells. At the same time, ALDH1A3 depletion increased the TGFB1 level in LNCaP cells, but had the opposite effect in the bone metastatic and androgen-independent cell lines C4-2B and PC3 (Figure 2A, Figure S1A). Also, we found that the knockdown of AR and ALDH1A1 in LNCaP cells was associated with inhibition of the TGF-β driven gene expression, whereas ALDH1A3 knockdown did not significantly affect gene targets of the TGF-β signaling (Figure 2B). Consistently, we observed a significant positive correlation of ALDH1A1 and a negative correlation of ALDH1A3 with TGFB1 in the PCa gene expression datasets TCGA PRAD (n = 493), MSKCC primary (n = 131), Broad/Cornell (n = 31), DKFZ (n = 118), and SMMU (n = 65) but not in the non-cancerous tissues (MSKCC cohort, n = 29) (Figure 2C).

**Figure 2.**
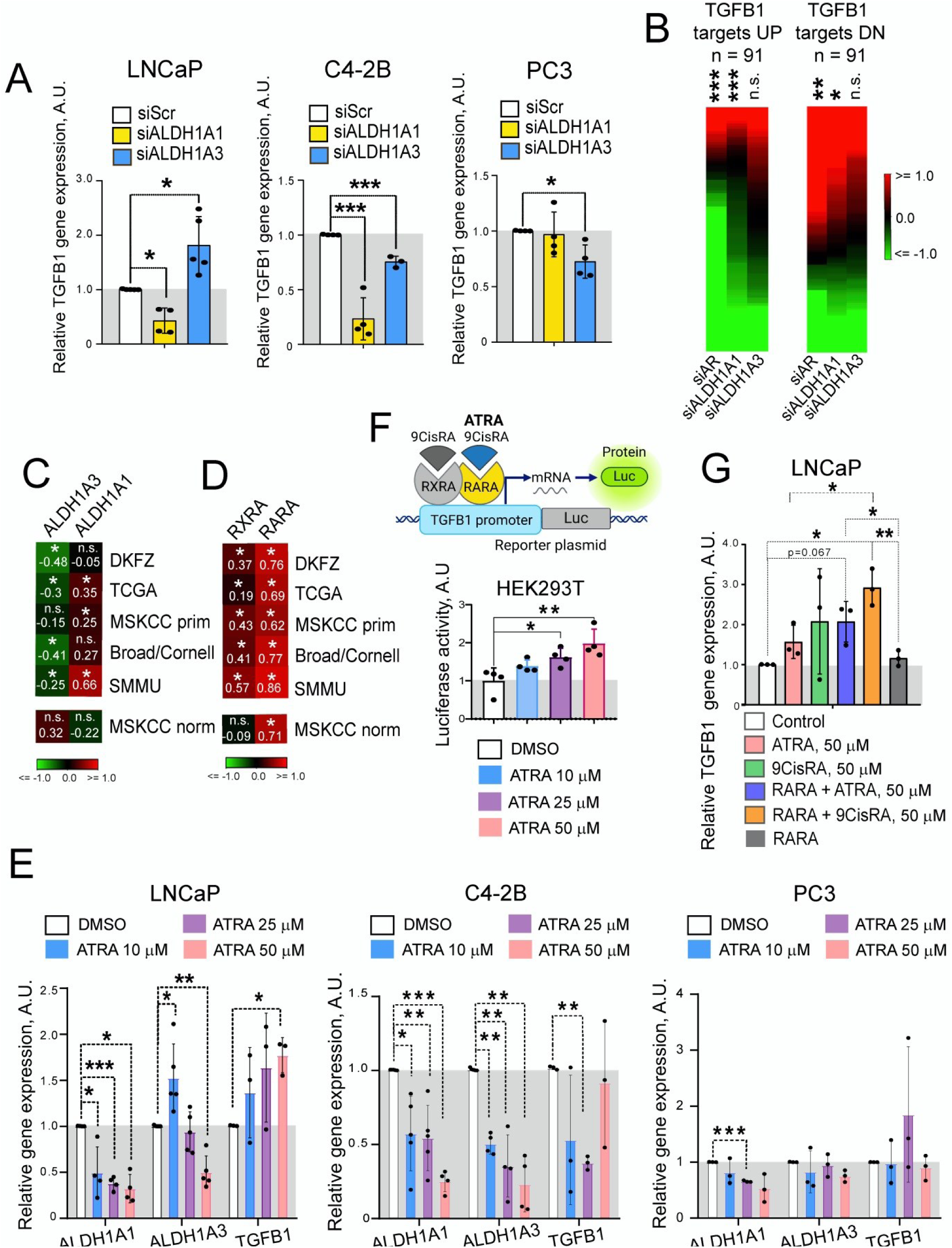
ALDH1A1 and ALDH1A3 regulate TGFB1 gene expression through RA-dependent mechanism. (**A**) Relative mRNA expression of TGFB1 gene in LNCaP, C4-2B and PC3 cells upon knockdown of ALDH1A1 or ALDH1A3 expression. Cells transfected with scrambled siRNA (siScr) were used as a control. n ≥ 3; Error bars = SD; *p < 0.05; ***p < 0.001. (**B**) Distribution of log2 fold change values (relative to siSCR control) for TGFB1 up or down genes. The gene sets for the TGFB1 responsive genes were described previously (48). Values are sorted from high to low in each column independently; therefore, the order of gene names is not the same. Significance of enrichment for up-or down-regulated genes evaluated by Wilcoxon signed rank test; *p < 0.05; **p<0.01; ***p < 0.001; n.s. – non-significant. (**C**) Correlation of ALDH1A1 and ALDH1A3 expression with TGFB1 mRNA levels in the PCa gene expression datasets TCGA PRAD (n = 493), MSKCC primary (n = 131), Broad/Cornell (n = 31), DKFZ (n = 118), and SMMU (n = 65) and non-cancerous tissues (MSKCC cohort, n = 29); *p < 0.05. (**D**) Correlation of RXRA and RARA expression with TGFB1 mRNA levels in the PCa gene expression datasets TCGA PRAD (n = 493), MSKCC primary (n = 131), Broad/Cornell (n = 31), DKFZ (n = 118), and SMMU (n = 65) and non-cancerous tissues (MSKCC cohort, n = 29); *p < 0.05. (**E**) LNCaP, C4-2B or PC3 cells were plated and treated with various concentrations of ATRA (10 μM, 25 μM, and 50 μM) or DMSO as a control for 48 h. Relative mRNA expression of ALDH1A1, ALDH1A3 and TGFB1 was measured by qPCR analysis. n ≥ 3; Error bars = SD; *p < 0.05; **p<0.01; ***p < 0.001. (**F**) HEK293T cells were transfected with pGL3-TGFB1 luciferase reporter plasmid (32) and treated with various concentrations of ATRA (10 μM, 25 μM, and 50 μM) and DMSO as a control. Luciferase activity was measured 48 h after the treatment started. n ≥ 3; Error bars = SD; *p < 0.05; **p<0.01. (**G**) qPCR analysis of TGFB1 expression in LNCaP cells upon transient RARA overexpression, treatment with ATRA or 9-cis retinoic acid (9CisRA) for 48 h or both. Cells transfected with empty plasmid were used as control. n = 3; Error bars = SD; *p < 0.05.

While ALDH1A1 and ALDH1A3 do not directly regulate gene expression, they play a role in synthesizing RA from retinol and therefore may potentially influence transcriptional programs driven by RA-dependent nuclear receptors such as RARA and RXRA. In the presence of RA, the transcription factors RARA and RXRA bind to RARE in target gene promoters, thereby modulating gene transcription. Analysis of TGFB1 expression in the above-described datasets TCGA, MSKCC primary, Broad/Cornell, DKFZ, and SMMU revealed its high correlation with the RARA and RXRA receptor in PCa but only with RARA in the non-cancerous tissues (Figure 2D). Therefore, we hypothesized that ALDH genes might regulate TGFB1 expression by RARA-dependent transcription. Next, we checked how the ATRA treatment affected TGFB1 mRNA levels. Cells were plated and treated with various concentrations of ATRA or DMSO as a control for 48 h. We observed a significant upregulation of the TGFB1 expression in response to the ATRA treatment in LNCaP cells but not in C4-2B and PC3 cells (Figure 2E). Treatment with ATRA inhibited the expression of ALDH1A1 and ALDH1A3 genes, consistent with our previous observations (29).

We further investigated the influence of ATRA on TGFB1 transcription using a luciferase reporter assay. To conduct a mechanistic analysis of the ATRA effect on TGFB1 expression, we used HEK293T cells possessing detectable basal levels of RARA and TGBF1 expression and highly amendable to transfection (Figure S2A). To investigate the ability of ATRA to regulate the transcription of TGFB1, HEK293T cells were transfected with pGL3-TGFB1 luciferase reporter plasmid described previously (32) and treated with various concentrations of ATRA and DMSO as a control. Luciferase activity was measured 48 h after the treatment started. We revealed an increased relative luciferase activity of the TGFB1 promoter upon ATRA treatment in a dose-dependent manner compared to control cells treated with DMSO (Figure 2F). Next, we transiently overexpressed RARA using previously described DNA construct (31) in PCa and LNCaP cells, treated them either with ATRA or 9CisRA, the products of ALDH1 biosynthesis having the highest affinity to RARs (21, 46, 47) and measured the mRNA expression of TGFB1 48 h after the treatment started. A combination of RARA overexpression and treatment with 50 μM of ATRA or 9CisRA resulted in more profound stimulation of TGFB1 gene expression than RARA overexpression or RA treatment alone (Figure 2G, Figure S2B). Notably, 9CisRA, a product specific for ALDH1A1 enzymatic activity, led to the higher TGFB1 gene expression activation compared to ATRA which is produced by both ALDH proteins (21, 22). These results demonstrate a signaling mechanism mediating the ALDH-dependent TGFB1 expression.

### RARA-and AR-interplay drives TGFB1 transcription

Next, we analyzed the impact of the RARA and AR levels on the TGFB1 expression by the knockdown experiments in the androgen-sensitive cell model. When AR was downregulated in LNCaP cells, the gene expression level of TGFB1 also decreased. For the RARA knockdown, a similar nearly significant trend was observed (Figure 3A). We analyzed the TGFB1 gene promoter by using the Eukaryotic Promoter Database and revealed putative RARE and AR-binding elements (Figure S2C). We, therefore, conducted a ChIP analysis in LNCaP cells with antibodies specific for AR and RARA proteins. The previously described RARA and AR transcription targets, RIG1 and KLK3, respectively, were used as positive controls (29). Coverage of all predicted binding sites was achieved by employing multiple primer pairs for each gene promoter. Cell pre-treatment with 50 μM of ATRA was used to induce RARA binding to RAREs in gene promoters (29). Our analysis revealed significantly increased precipitation of different promoter regions of TGFB1 with RARA and AR antibodies compared to the control IgG antibody (Figure 3B). These results suggest that TGFB1 is regulated by ALDH1A1 and ALDH1A3 genes in RAR-and AR-dependent manner.

**Figure 3.**
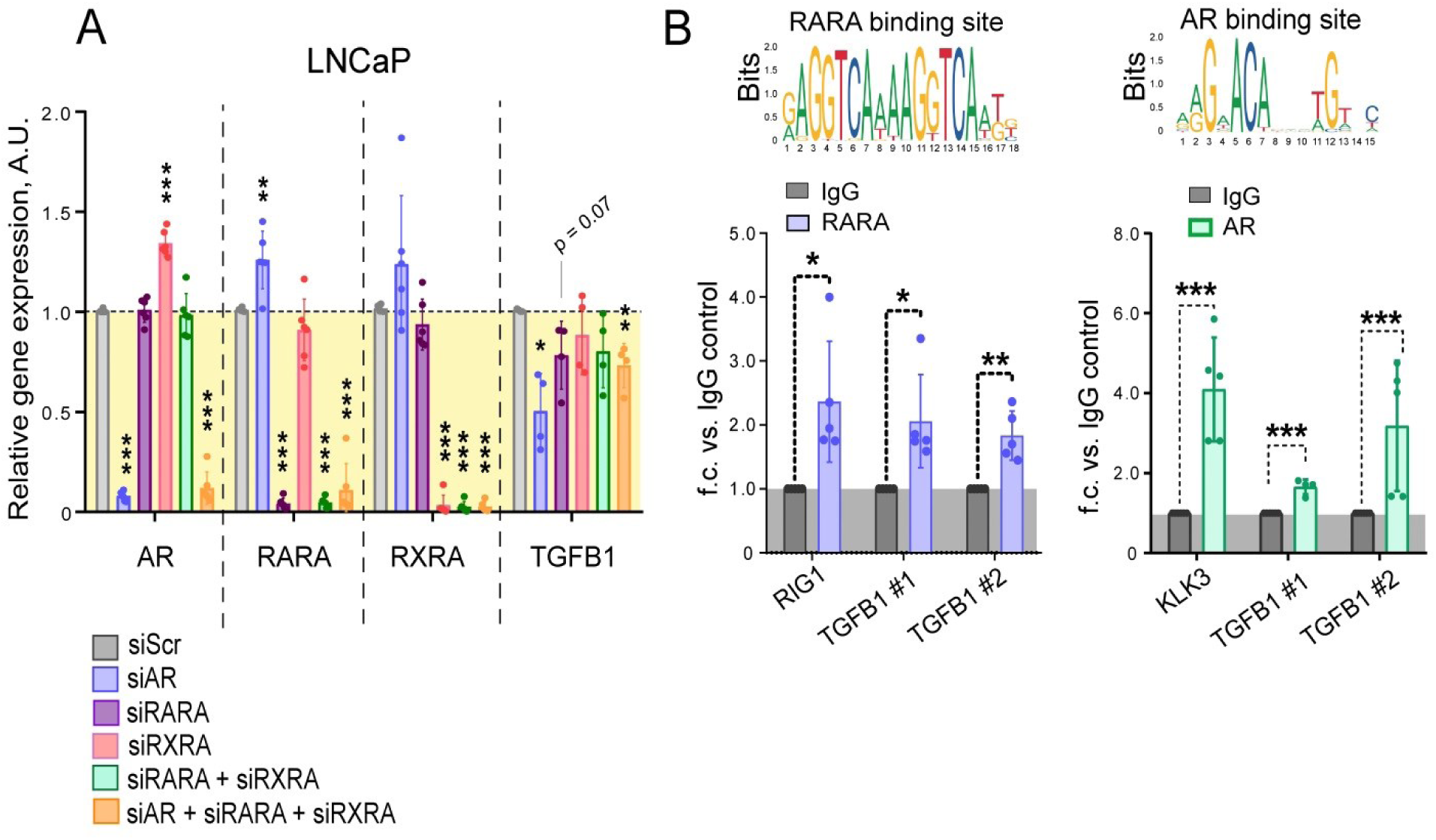
RARA-and AR-interplay drives TGFB1 transcription. (**A**) qPCR analysis of the relative TGFB1 expression in LNCaP cells upon knockdown of AR, RARA, and RXRA expression alone or in combination. Cells transfected with scrambled siRNA (siScr) were used as a control. n = 3; Error bars = SD; *p < 0.05; **p < 0.01; ***p < 0.001. (**B**) Chromatin immunoprecipitation (ChIP)-qPCR analysis in LNCaP cells for assessment of the direct binding of RARA and AR proteins to the putative binding regions in the promoter of the TGFB1 gene. RARA and AR binding sites were taken from the JASPAR CORE database (49). n ≥ 3; Error bars = SD; *p < 0.05; **p < 0.001; ***p < 0.001.

### TGF-β1 regulates members of the MMP family

TGF-β1-dependent signaling also contributes to the metastatic process by regulating the MMPs’ expression (9, 50). MMPs are enzymes involved in the degradation of the ECM (8). They play an important role in cancer-associated bone remodeling, thus facilitating intraosseous tumor spread (51). We used the ROC curve analysis to assess the potential association of the expression levels of 20 MMP genes with BRFS in the TCGA PRAD gene expression dataset. This analysis revealed that only MMP11 and MMP26 gene expression levels are significantly but oppositely associated with clinical outcomes (Figure 4A). Notably, MMP11 was previously described as one of the TGF-β1-responsive genes (52). Indeed, we observed a significant positive correlation of TGFB1 with MMP11 and a significant negative correlation with MMP26 expression in several PCa gene expression datasets, including primary PRAD TCGA (38) (n = 493) cohort, MSKCC primary (40) (n = 131), Broad/Cornell (42) (n = 31), SMMU (41) (n = 65) and DKFZ (41) (n = 118), but not in the non-cancerous tissues (MSKCC cohort, n = 29) (Figure 4B).

**Figure 4.**
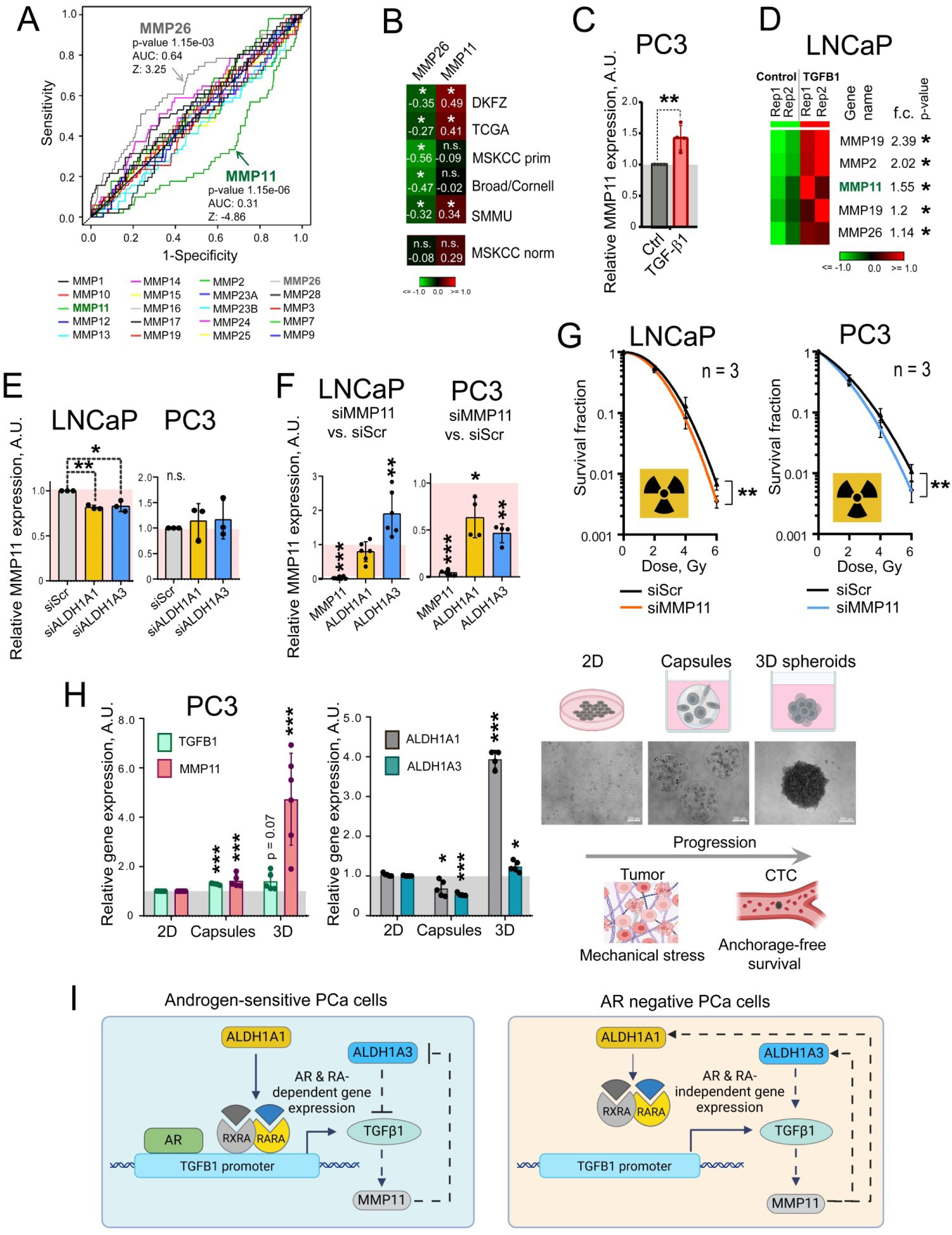
TGFβ1 regulates the members of the MMP family. (**A**) The ROC curve analysis of the potential association of the expression levels of 20 MMP genes with biochemical recurrence-free survival (BRFS) in the TCGA PRAD gene expression dataset (n = 493). ROC analysis was conducted using easyROC web-tool (44). (**B**) Correlation of MMP11 and MMP26 gene expression with TGFB1 mRNA levels in the PCa gene expression datasets TCGA PRAD (n = 493), MSKCC primary (n = 131), Broad/Cornell (n = 31), DKFZ (n = 118), and SMMU (n = 65) and non-cancerous tissues (MSKCC cohort, n = 29); *p < 0.05. (**C**) PC3 cells were treated with 5 ng/ml of TGFβ1 for 48 h, and MMP11 levels were analyzed by qPCR. n = 3; Error bars = SD; **p < 0.01. (**D**) Analysis of the previously published dataset (54) indicated that TGF-β signaling in stroma cells induces MMP expression in LNCaP cells. LNCaP cells overexpressing TGFB1 and co-cultured with stroma cells were compared to the control LNCaP co-cultured with stroma. (**E**) qPCR analysis of the relative MMP11 expression in LNCaP and PC3 cells upon knockdown of ALDH1A1 or ALDH1A3. Cells transfected with scrambled siRNA (siScr) were used as controls. n = 3; Error bars = SD; *p < 0.05; **p < 0.01. (**F**) qPCR analysis of the relative MMP11, ALDH1A1 and ALDH1A3 expression in LNCaP and PC3 cells upon MMP11 knockdown. Cells transfected with scrambled siRNA (siScr) were used as controls. n = 3; Error bars = SD; *p < 0.05; **p < 0.01; ***p < 0.001. (**G**) Relative cell radiosensitivity was analyzed by 2D radiobiological colony forming assay after siRNA-mediated knockdown of MMP11 in LNCaP and DU145 cells. Cells transfected with scrambled (Scr) siRNA were used as controls. N≥3; Error bars = SD; **p<0.01. (**H**) PC3 cells were cultured either under 2D conventional adherent conditions, in the polymer-based microcapsules mimicking mechanical stress within tumors (30), or in 3D anchorage-free cultures as a model of the intermediate stage of metastasis, circulating tumor cells (CTC). In all conditions, cells were kept in the same culture media. At day 3, the cells were collected, and relative gene expression of TGFB1, MMP11, ALDHA1 and ALDHA3 was analyzed by qPCR. n ≥ 3; Error bars = SD; *p < 0.05; ***p < 0.001. (**I**) ALDH1A1 regulates TGFB1 expression in androgen-sensitive cells through AR-and RA-dependent mechanisms. In contrast, the interplay between TGF-β1 and MMP11 is present in the androgen-sensitive and castration-resistant models of PCa (like in LNCaP and PC3 cells, respectively).

Next, we analyzed how the TGF-β1 ligand affects the levels of MMP gene expression. Since LNCaP cells have a mutated TGF-β receptor I (TβRI/ALK-5), making them insensitive to TGF-β1 treatment (53), we have used PC3 as a TGF-β1 responsive PCa model. Treatment of PC3 cells with 5 ng/ml of TGFβ1 for 48 h upregulated MMP11 expression, whereas the levels of MMP26 mRNA expression were below the levels of qPCR detection (Figure 4C). Of note, analysis of a previously published gene expression dataset confirmed the upregulation of several MMPs, including MMP11 and MMP26, in the co-culture of prostate stroma and LNCaP cells overexpressing TGFB1 compared to the control LNCaP and stroma co-culture indicating that prostate stromal cell-specific TGF-β signaling induces MMP11 expression in LNCaP cells (54) (Figure 4D). The gene expression level of MMP11 is decreased after ALDH1A1 and ALDH1A3 knockdown in the androgen-sensitive LNCaP cells, but not in the AR-negative PC3 cells (Figure 4E, Figure S1A). The feedback loop mechanisms also differ in these cell models: the knockdown of MMP11 induces ALDH1A3 expression in LNCaP cells and inhibits ALDH1A1 and ALDH1A3 expression in PC3 cells (Figure 4F). Increased cell radioresistance is one of the features of prostate CSCs (29). Knockdown of MMP11 increased cell radiosensitivity in LNCaP and PC3 cells (Figure 4G, Figure S2DE). Additionally, we have cultured PC3 cells in anchorage-independent spheroid cultures or polymer-based microcapsules to generate stiff microenvironment (30). Gene expression levels in 3D conditions were compared to the 2D conditions. In all conditions, cells were kept in the same culture media. Expression levels of ALDH1A1 and ALDH1A3 were used as CSC markers. We could show that 3D culture conditions mimicking mechanical stress and anchorage-free conditions induced the upregulation of MMP11 and TGFB1 expression (Figure 4H). Altogether, this data suggests that ALDH1A1 positively regulates TGFB1 expression in the RARA-and AR-dependent manner in the androgen-sensitive PCa cells, however, the interplay between TGF-β1 and MMP11 levels could be present at the androgen-sensitive and castration-resistant stages of PCa (Figure 4I). Both genes are upregulated by the cellular stresses encountered by tumor cells during tumor progression and metastatic dissemination.

### Gene expression levels of MMP11 as a potential predictor of PCa patients’ outcome

The potential role of MMP11 and MMP26 as prognostic biomarkers was first analyzed in the publicly available gene expression datasets. Among several analyzed clinical parameters, the level of MMP11 expression correlated the most with the Gleason score in several independent datasets (PRAD TCGA (38), n = 498; DKFZ (41), n = 118; and MSKCC primary and metastatic (40), n = 150) (Figure 5A, Figure S3A). In contrast, MMP26 level were negatively associated with the Gleason score in the PRAD TCGA dataset and had no significant association with the Gleason score in other analysed cohorts (Figure 5B, Figure S3B).

**Figure 5.**
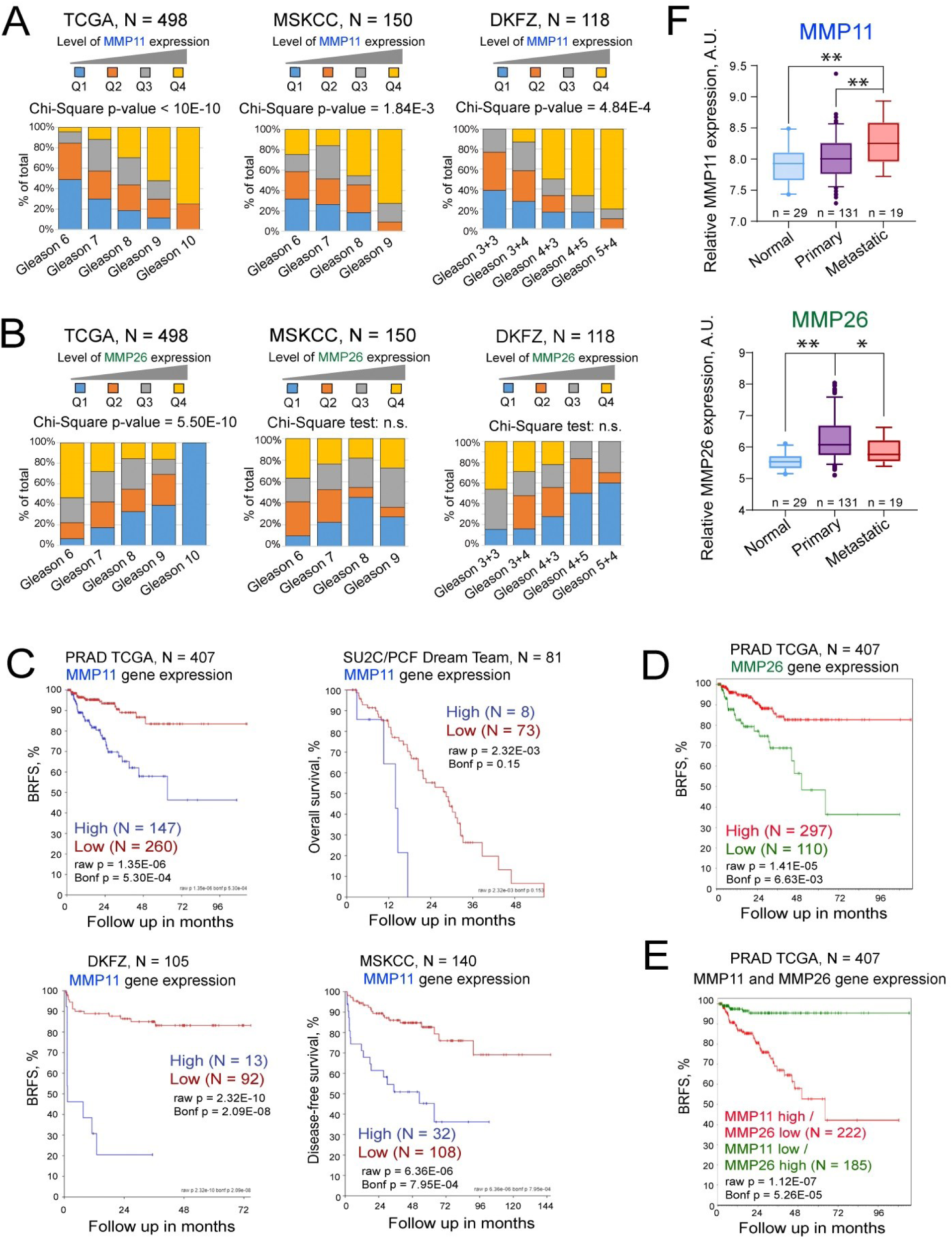
Gene expression levels of MMP11 as a potential predictor of PCa patients’ outcome. (**A**) MMP11 and (**B**) MMP26 mRNA expression levels were correlated with Gleason score in several independent datasets (PRAD TCGA, n = 498; DKFZ, n = 118; and MSKCC primary and metastatic, n = 150). Q: quartile. (**C**) The Kaplan-Meier analyses of the association of MMP11 expression and clinical outcomes in the PCa gene expression datasets TCGA PRAD (n = 407), MSKCC (n = 140), DKFZ (n = 105), and SU2C/PCF Dream Team (n = 81). The patients were stratified by the most significant cut-off for MMP11 expression levels. (**D**) The Kaplan-Meier analyses of the association of MMP26 expression and BRFS in the TCGA PRAD PCa gene expression dataset (n = 407). The patients were stratified by the most significant cut-off for MMP26 expression levels. (**E**) A combined MMP11 high/MMP26 low gene expression signature improves the prediction of BRFS in the TCGA PRAD PCa gene expression dataset (n = 407). (**F**) A relative expression of MMP11 and MMP26 genes in non-cancerous tissues, primary and metastatic tumors (MSKCC dataset). Statistics was performed by one-way ANOVA followed by posthoc Tukey HSD test. The error bars denote the 5th and 95th percentile values. *p < 0.05; **p < 0.01.

Analysis of the PRAD TCGA (38) cohort (n = 407) revealed that MMP11 and MMP26 expression levels have an opposite and significant association with BRFS, and MMP11 high / MMP26 low gene expression signature even further improves the outcome prediction (Figure 5CDE). We also were able to validate our finding on other independent cohorts, demonstrating a positive association of high MMP11 levels with shorter OS (SU2C/PCF Dream Team, n = 81); DFS (MSKCC primary (40), n = 140) and BRFS (DKFZ (41), n = 105) (Figure 5C). Only one cohort, based on the analysis of the FFPE tissues (Manchester dataset, n = 432, Table 3), showed the opposite correlation Figure S4A). This discordance can be explained by the different stability of the RNA in the frozen and FFPE tissues, which can be critical for low-abundant genes such as MMPs (55) (Figure S4B). The role of MMP11 in tumor development is further supported by its increased expression in PCa compared to normal tissues and even higher expression in metastases in the MSKCC (40) dataset (n = 179). In contrast, MMP26 expression was significantly downregulated in metastases compared to primary tumor tissues (Figure 5F). These findings suggest that MMP11 gene expression levels are upregulated during tumor progression and could be used as a potential predictor of PCa patients’ outcome.

### The protein MMP11 level is a prognostic plasma-based biomarker for patients with metastatic PCa

To further validate our findings, we performed an ELISA-based analysis of MMP11 and TGF-β1 levels in plasma samples from the independent patient cohorts (Figure 6A). First, we analyzed samples from patients with oligometastatic PCa treated with local ablative radiotherapy (n = 31) (Table 1) and healthy donors (n = 5) and found significantly higher concentrations of MMP11 in the samples from PCa patients, whereas TGF-β1 has shown a similar but not significant trend (Figure 6B). Next, we analyzed a potential association between plasma MMP11 and PSA increase in these patients and detected a significant positive association between high MMP11 levels and rapid biochemical progression (Figure 6C, Figure S5A).

**Figure 6.**
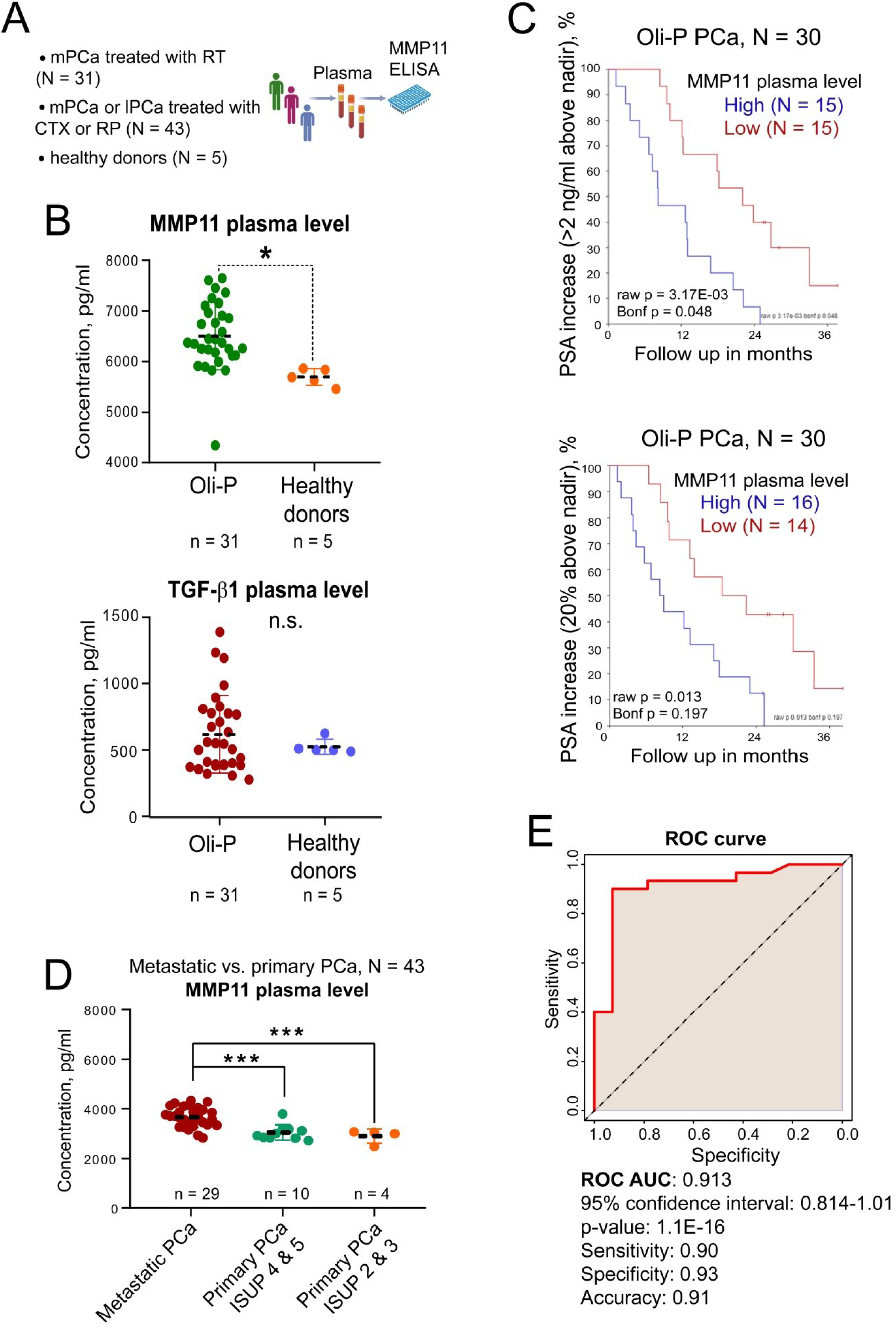
The protein MMP11 level is a prognostic liquid biopsy-based biomarker. (**A**) Scheme of the MMP11 ELISA of plasma samples; omPCa: oligometastatic prostate cancer; mPCa: metastatic prostate cancer; lPCa: local prostate cancer; RT: radiotherapy; CTX: taxane-based chemotherapy; RP: radical prostatectomy. (**B**) ELISA analysis of the MMP11 protein levels in the plasma samples (n = 31) from patients with oligometastatic PCa treated with local ablative radiotherapy at the Department of Radiotherapy and Radiation Oncology of the University Hospital Carl Gustav Carus, Dresden (ClinicalTrials.gov Identifier: NCT02264379) and healthy donors (n = 5); *p < 0.05. (**C**) The Kaplan-Meier analyses of the association of MMP11 plasma levels and PSA increase (>2 ng/ml above nadir and 20% above nadir) in patients with oligometastatic PCa treated with local ablative radiotherapy, n = 30. (**D**) ELISA analysis of the MMP11 protein levels in plasma samples (n = 43) from patients with localized or metastatic PCa (mPCa) treated with radical prostatectomy (primary PCa) or taxane-based chemotherapy (mPCa), respectively, at the Department of Urology of the University Hospital Carl Gustav Carus, Dresden. ISUP - the International Society of Urological Pathology grade. (**E**) The ROC curve analysis of the discriminatory power of plasma MMP11 levels as a potential blood-based biomarker of metastases. ROC analysis was conducted using ROC Plotter web-tool (43).

We have additionally validated MMP11 as a marker of metastases using plasma samples (n = 43) from patients with localized or metastatic PCa (hormone-sensitive PCa or CRPC) treated with radical prostatectomy (primary PCa) or taxane-based chemotherapy, docetaxel or cabazitaxel (metastatic PCa) (Table 2). Out of 29 patients with metastatic PCa, 27 patients (93%) have bone metastases. We observed significantly higher concentrations of MMP11 in the samples from PCa patients with metastatic disease compared to the samples from patients with localized PCa (Figure 6D).

The discriminative power of serum MMP11 level as a potential blood-based biomarker of metastases was analyzed using the ROC curve analysis (Figure 6E, Figure S5B), which revealed a high specificity (0.93) and sensitivity (0.9) of the biomarker (determined by Youden index). The AUC value was 0.91, indicating a good biomarker potential to differentiate between metastatic and non-metastatic PCa. All these findings suggest that patients with pre-treated and highly advanced PCa have higher MMP11 plasma levels than patients with localized PCa. Furthermore, plasma levels of MMP11 are a promising prognostic biomarker in patients with androgen-sensitive oligometastatic PCa receiving ablative EBRT to the metastatic lesions.

Furthermore, analysis of the MMP11 expression in the broad panel of tumor samples and normal tissues using the GEPIA 2 web tool (56) suggested that in addition to PCa, it is overexpressed in almost all types of cancer (Figure S5C). As a proof of concept study, we analyzed the MMP11 plasma levels in the NMRI (nu/nu) tumor-free healthy mice and in the NMRI (nu/nu) mice bearing subcutaneous head and neck squamous cell carcinoma (HNSCC) xenograft tumors and found a statistical trend toward increased MMP11 levels in the tumor-bearing mice (Figure S5D). This finding suggests that MMP11 could serve as a plasma-based biomarker for some other types of cancer.

### Proteomic profiling of plasma samples

Based on the significant correlation of MMP11 levels with PSA increase, we have selected for proteomics profiling the plasma samples from patients with short PSA-free survival and high MMP11 levels (n=5) and patients with extended PSA-free survival and low MMP11 levels (n=5) (Figure 7A-E); samples were analyzed using an LC-MS/MS label-free approach (57). In total, the set of 344 plasma proteins was identified as listed in Table S3, among which 297 proteins were assigned to the PANTHER protein class (58), 199 proteins were mapped to the GO molecular function terms, and 73 proteins were associated with pathways according to the PANTHER classification. Figure S6A-C shows a functional classification of the identified proteins, their molecular functions according to PANTHER GO-slim, and associated pathways. The defense/immunity proteins (PC00090, n *=* 82) represent the most abundant protein class, followed by the protein-binding activity modulators (PC00095, n *=* 33), metabolite interconversion enzymes (PC00262, n *=* 31), protein modifying enzymes (PC00260, n *=* 30), cytoskeletal proteins (PC00085, n *=* 28) and transfer/carrier proteins (PC00219, n *=* 23).

**Figure 7.**
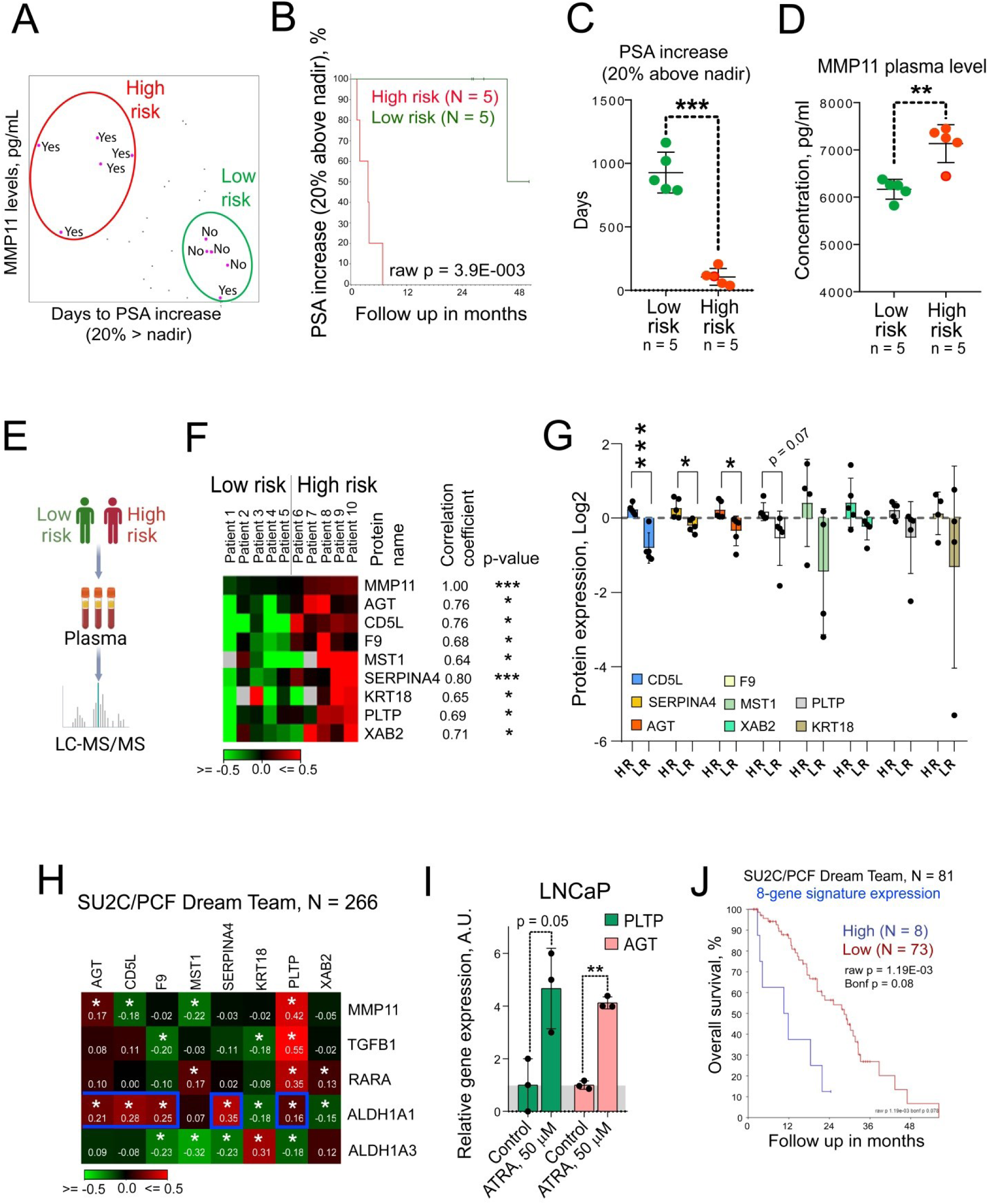
Development of the ALDH1A1/MMP11-related plasma proteome-based prognostic signature. (**A**) “High-risk” and “low-risk” samples were selected in the Oli-P study cohort based on PSA-free survival time and MMP11 levels measured by ELISA assay. “Yes” or “no” indicate the occurrence or absence of PSA increase (20% above nadir), correspondingly. (**B**) The Kaplan-Meier analysis of the PSA increase in high risk (N = 5) and low risk group (N = 5). (**C**) Days to PSA increase in high risk (N = 5) and low risk group (N = 5); Error bars = SD; ***p < 0.001. (**D**) MMP11 levels measured by ELISA assay in high risk (N = 5) and low risk group (N = 5); Error bars = SD; **p < 0.01. (**E**) Schematic diagram of the proteomics analysis of plasma samples. (**F**) A subset of 8 non-immunoglobulin proteins detected by mass spectrometry that significantly correlate with MMP11 ELISA values; ***p < 0.001; *p < 0.05. (**G**) Abundances of 8 selected plasma proteins in high-risk (HR; N = 5) and low-risk group (LR; N = 5); Error bars = SD; ***p < 0.001; *p < 0.05. (**H**) Analysis of the potential correlation between the expression levels of 8 genes coding selected plasma proteins and 5 genes from ALDH1/MMP11 pathway in the SU2C/PCF Dream Team cohort of the patients with metastatic PCa (N = 266); *p < 0.05. (**I**) LNCaP cells were treated with ATRA at a concentration of 50 μM or DMSO as a control for 48 h. Relative gene expression was analyzed using RNA sequencing analysis as described previously (29). n ≥ 3; Error bars = SD; *p < 0.05; **p<0.01; ***p < 0.001. (**J**) The Kaplan-Meier analyses of the association of 8-gene signature expression and clinical outcomes in the PCa gene expression dataset SU2C/PCF Dream Team (n = 81). The patients were stratified by the most significant cut-off for gene expression levels.

From the whole proteomic dataset, 8 non-immunoglobulin proteins with the highest positive correlation with MMP11 were selected: SERPINA4, AGT, CD5L, F9, MST1, PLPT, XAB2 and KRT18 (Figure 7F). In this subset, 3 proteins (SERPINA4, AGT, and CD5L) were significantly upregulated in the high-risk patient cohort, whereas 4 proteins (F9, MST1, PLPT, and XAB2) have shown a similar statistical trend (Figure 7G). Moreover, we analyzed the expression of the corresponding genes in the SU2C/PCF Dream Team cohort of the patients with metastatic PCa (39). Five of the corresponding genes, PLTP, SERPINA4, AGT, CD5L, and F9 possess a significant positive correlation with the ALDH1A1 gene expression, and two of them (PLTP and AGT) maintain a significant positive correlation with the MMP11 gene expression (Figure 7H). Consistent with this observation, both PLTP and AGT genes are upregulated in response to ATRA treatment (Figure 7I), suggesting that the ALDH enzymatic products can directly regulate their transcription. Furthermore, a signature combining the expression of all 8 genes has a significant correlation with overall survival in the SU2C/PCF Dream Team cohort of patients with metastatic PCa (Figure 7J). Therefore, a signature of plasma proteins linked to the ALDH1A1/MMP11 axis appeared as a promising prognostic tool for patients with PCa.

## Discussion

Metastatic dissemination is a complex evolutionary process that includes multiple steps in selecting tumor cells for their ability to escape from the primary tumor, survive in the bloodstream, disseminate to distant organs, and initiate secondary tumor growth. The metastasis-initiating cells (MICs) possess the critical features of CSCs to self-renew and to give rise to more differentiated progenies. The functional and phenotypical similarities between MICs and CSCs suggest that tumor metastases are driven by the subpopulations of CSCs, which evolved or induced during tumor progression and metastatic cascade (45, 59). Metastatic PCa has unfavorable outcomes, and the therapeutic responses of patients are very heterogeneous. The limited therapeutic options for patients with metastatic PCa and the lack of prognostic biomarkers of metastatic disease are attributed to the poor understanding of the mechanisms mediating metastatic dissemination. Thus, identifying the metastasis mechanisms is of utmost importance in developing new biomarkers for metastatic prognosis and therapeutic targets for metastatic prevention in PCa.

In our study, we demonstrated for the first time the interplay between stem cell regulating genes ALDH1A1 and ALDH1A3 and the metastasis-driven signaling mechanism mediated by TGF-β1 and MMP11. We demonstrated that TGFB1 mRNA transcription is directly regulated by ALDH1A1 and ALDH1A3 genes in RAR-and AR-dependent manner. We confirmed previous observations that MMP11 is one of the TGF-β1-regulated genes and analyzed the role of MMP11 as a potential plasma biomarker. To our knowledge, this is the first report revealing that MMP11 is a promising liquid biopsy-based prognostic biomarker for patients with metastatic PCa.

Until recently, high enzymatic activity of ALDH proteins in the populations of tumor-initiating cells has been considered a correlative marker of prostate CSCs that does not affect stem cell properties. The evolving understanding of CSC biology has reshaped a view of ALDH activity, shifting it from a mere marker to a key regulator of stemness across various tumor types, including PCa (19). ALDH metabolic enzymes primarily function by oxidizing cellular aldehydes to carboxylic acids in a NAD(P)+ dependent manner, concurrently generating NAD(P)H. These resultant products play pivotal roles in maintaining cellular homeostasis and promoting survival. Notably, certain carboxylic acids produced through ALDH-mediated reactions, such as the neurotransmitter γ-aminobutyric acid (GABA), and RA isomers like ATRA and 9CisRA, act as ligands for nuclear receptors such as RARs and RXRs (21, 46, 47). Treatment with ATRA is a potent differentiation therapy for acute promyelocytic leukemia (APL) (23). Therefore, RA-dependent signaling has been long considered solely as an inducer of normal stem cells and CSC differentiation and apoptosis (60, 61). Nevertheless, recent findings by our team and others suggest that ALDH-driven RAR/RXR-mediated transcription of DNA repair and metastatic genes plays a vital role in therapy resistance and metastatic spread (29, 62, 63). Furthermore, cell response to the products of the ALDH biosynthesis depends on the type of synthesized RA isomers. Various ALDH-produced RA ligands bind to the members of the RAR/RXR family with different affinity and, correspondingly, induce distinct transcriptional programs (64–66). Both ATRA and 9CisRA are potent activators of RARs, whereas RXRs have a high affinity for 9CisRA (65). RARs act as transcriptional regulators when they are forming heterodimers with RXRs, while RXRs can activate transcription as homodimers. Upon binding to retinoid ligands, RXR/RAR dimers recognize specific DNA sequences known as RAREs with a high affinity, thereby modulating the transcription of target genes. Apart from RARs, RXR nuclear receptors interact with other transcriptional regulators such as peroxisome proliferator-activated receptors (PPARs), liver X receptors (LXRs), farnesoid X receptor (FXR), and other transcriptional factors involved in the metabolic and developmental pathways (64). Of importance, RARs and RXRs are reported to regulate AR target genes, and the interplay between AR and RAR/RXR is essential for the governance of prostate-specific gene expression (67).

This study demonstrated that both RARA and AR positively affect the expression of TGFB1 through direct binding to its promoter, as shown in Figure 4G. The products of ALDH catalytic activity ATRA or 9CisRA induce expression of TGFB1, whereas 9CisRA, mainly produced by ALDH1A1, had a more profound effect on TGFB1 gene expression compared to ATRA produced by both ALDH proteins. This ligand-dependent effect on gene transcription can explain the disparate impact of ALDH1A1 and ALDH1A3 on global gene expression (29) and, consequently, their distinct role in PCa progression. Indeed, we showed that the two ALDH genes differently regulate *in vitro* cell migration and TGFB1 gene expression, and the mode of this regulation depends on the androgen sensitivity in the used PCa models. Our findings suggest that ALDH1A1 positively and ALDH1A3 negatively regulate *in vitro* cell migration and TGFB1 expression in the AR-positive LNCaP and C4-2B cells. Consistent with *in vitro* data and the results of our recent studies (29), we also found that ALDH1A1 and ALDH1A3 are differently associated with BRFS and MFS in patients with high-risk locally advanced PCa treated with ADT, and high ALDH1A1 expression is significantly associated with worse outcomes. We confirmed a significant positive correlation of TGFB1 and ALDH1A1, RARA, and RXRA gene expression and a negative correlation of TGFB1 and ALDH1A3 in several PCa patient gene expression datasets, suggesting that ALDH1A1-driven and RAR/RXR-mediated TGFB1 expression can be clinically relevant.

TGF-β1 plays a dual role in PCa development. At the early stages of cancer progression, it acts as a tumor suppressor, whereas at later stages, it promotes tumor formation by stimulating proliferation, invasion, and metastasis (12, 68). The elevated preoperative plasma levels of TGF-β1 in patients undergoing radical prostatectomy were associated with increased risk of lymph node and skeletal metastases and further PCa progression (69, 70). Once cancer spreads to the bones, TGFB1 plays a pleiotropic role in the regulation of the tumor microenvironment by stimulating PCa cell proliferation and migration and inducing bone remodeling and angiogenesis (71).

Regulation of the MMP expression is one of the key mechanisms mediating TGF-β1-dependent PCa invasion (9, 10). MMPs are a large family of endopeptidases disrupting the basement membrane and ECM components (8). We demonstrated that among 20 members of the MMP family analyzed in our study, only MMP11 and MMP26 gene expression levels are significantly (but oppositely) associated with PCa patients’ clinical outcomes as was also confirmed by other authors (72–74). In line with these observations, we found a positive correlation of TGFB1 with MMP11 and a negative correlation with MMP26 expression in several analyzed PCa patient gene expression datasets. We confirmed that MMP11 is positively regulated by TGF-β1 *in vitro* and associated with unfavorable clinical outcomes in several independent PCa cohorts. Within the plasma of oligo-progressive PCa patients, we detected a significantly higher MMP11 concentration compared to the healthy donors. In our patient cohort, the high level of MMP11 in the plasma was associated with a faster biochemical relapse. Our study suggests that plasma MMP11 has a promising biomarker potential for metastatic PCa. Furthermore, we have proposed the ALDH1A1/MMP11-related plasma proteome-based prognostic signature, laying a solid ground for future studies.

This study’s limitations include a small cohort size for analyzing MMP11 plasma levels and proteomic profiling, as well as its retrospective character. Further independent validation of MMP11 and related proteins as plasma biomarkers in the prospective study for the larger patient cohort is warranted to confirm their role as a clinical prognosticator and potential therapeutic target for PCa treatment. MMP11 targeting in other types of cancer has already been identified as a potential therapeutic strategy due to its role in tumor progression and metastasis. MMP11-directed vaccine induced cell-mediated and antibody immune response and exerted significant antitumor protection in mice with colon cancer in prophylactic and therapeutic settings (75). Furthermore, previous research has demonstrated that androgen-induced miR-135a acts as a tumor suppressor in PCa cells by downregulating MMP11, and this mechanism was associated with developing androgen resistance. These findings suggest an additional route of MMP11 regulation in androgen-dependent PCa and its role in PCa progression to the CRPC stage (76).

## Conclusions

This report provides the first evidence that MMP11 plasma levels is higher in patients with pre-treated and highly advanced PCa than in patients with localized PCa. Furthermore, plasma levels of MMP11 are a promising prognostic biomarker in patients with androgen-sensitive oligometastatic PCa receiving ablative radiotherapy for metastatic lesions. MMP11 could potentially serve as a prognostic biomarker for patients with aggressive PCa and a provisional target to eliminate metastasis-initiating cell populations. Moreover, a hypothetical ALDH1A1/MMP11-related plasma proteome-based prognostic signature was identified, paving the way for further clinical studies.

## List of abbreviations

ADT: androgen deprivation therapy
ALDH: aldehyde dehydrogenase
AR: androgen receptor (AR)
aRT: ablative radiotherapy
AUC: area under the ROC curve
BRFS: biochemical recurrence free survival
CAPRA: Cancer of the Prostate Risk Assessment
ChIP: chromatin immunoprecipitation
CRPC: castration resistant prostate cancer
CTC: circulating tumor cells
DFS: disease free survival
EBRT: external beam radiotherapy
EMT: epithelial mesenchymal transition
HDR: high dose rate
ISUP: International Society of Urological Pathology
GTV: gross tumor volume
ELISA: enzyme-linked immunosorbent assay
MFS: metastatic free survival
MMP11: matrix metalloproteinase 11
NCCN: National Comprehensive Cancer Network
OS: overall survival
RA: retinoic acid
PCa: prostate cancer
PLK3: polo-like kinase 3
PSA: prostate specific antigen
RARA: retinoic acid receptor alpha
ROC: receiver operating characteristic
RXRA: Retinoid X receptor alpha
TGFB1: transforming growth factor-beta 1

## Declarations

### Consent for publication

Not applicable

### Availability of data and materials

The datasets used and analyzed during the current study are available from the corresponding author on reasonable request

### Author Contributions

IG, VL, and AD conceived the study. IG, VL, MG, MS, JP, ASK, UK performed the experiments; IG, VL, MS, DK, CP, FL, TH, YA and MG acquired the data; IG, VL, AL, FL, TH, KE, SF, AB, MR, AC, MP and AD analyzed the data; AD supervised the experiments; AD and IG wrote the manuscript; VL, IG, MR, AC, ALE, RKW, TH, CP, CT, MG, PW, MP, MK and AD edited the manuscript; AD supervised the study and provided project administration. All the authors reviewed the final version.

## Acknowledgments

We appreciate Andrea Petzold for technical assistance with handling of the blood samples.

## Supplementary information

**Table S1.**
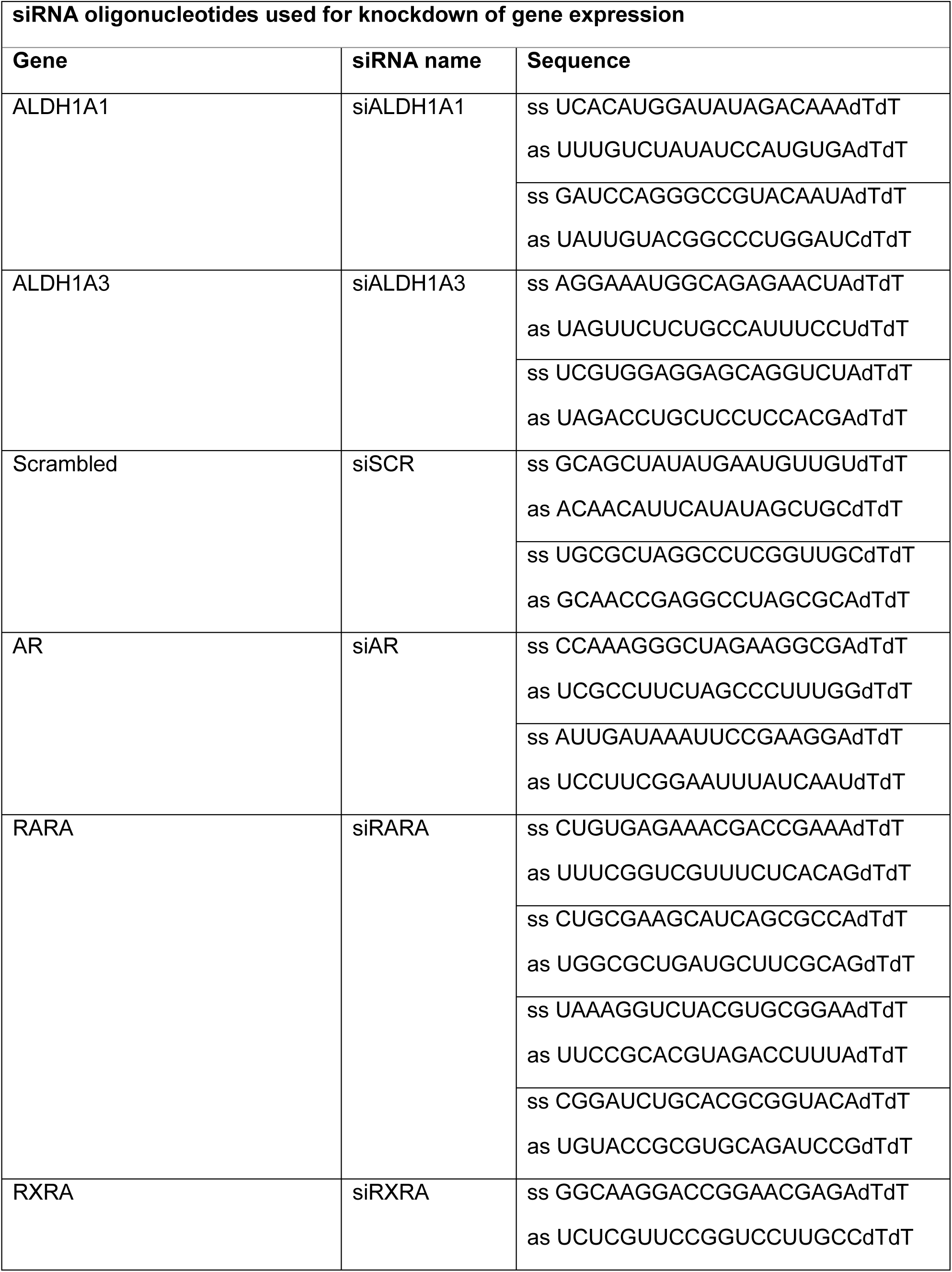
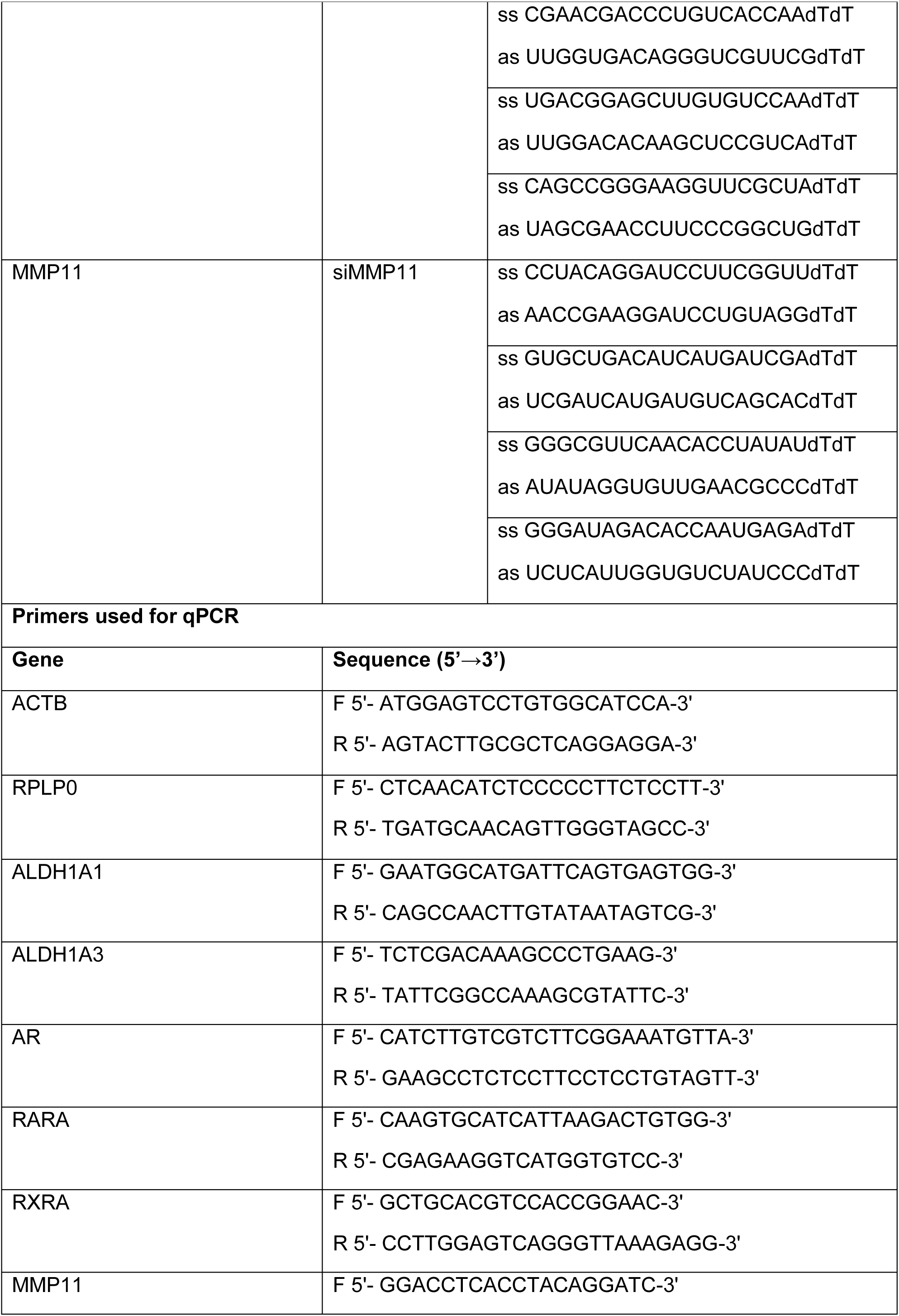
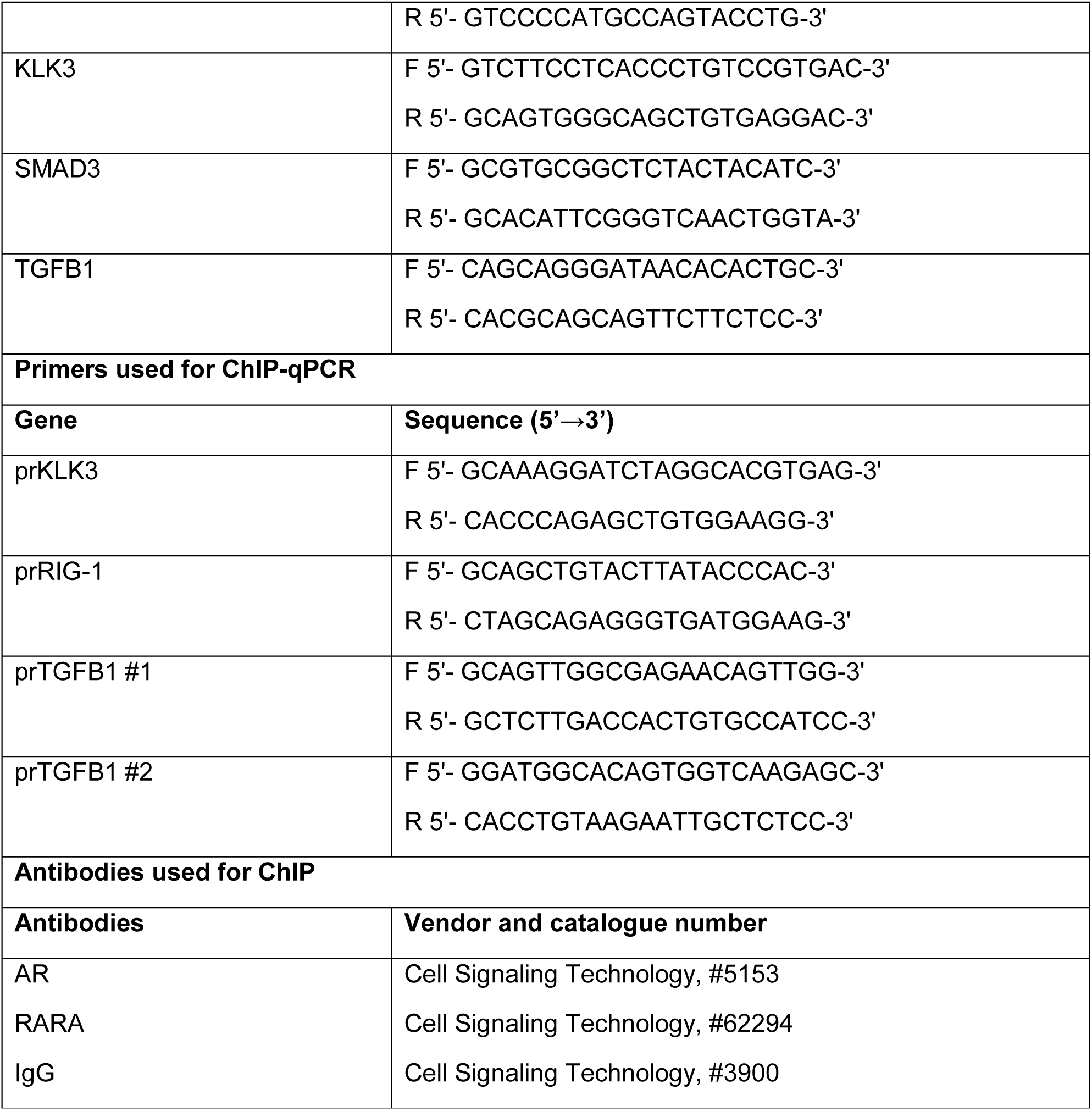
Primers, and siRNA oligonucleotides and antibodies used for the study.

**Table S2.**
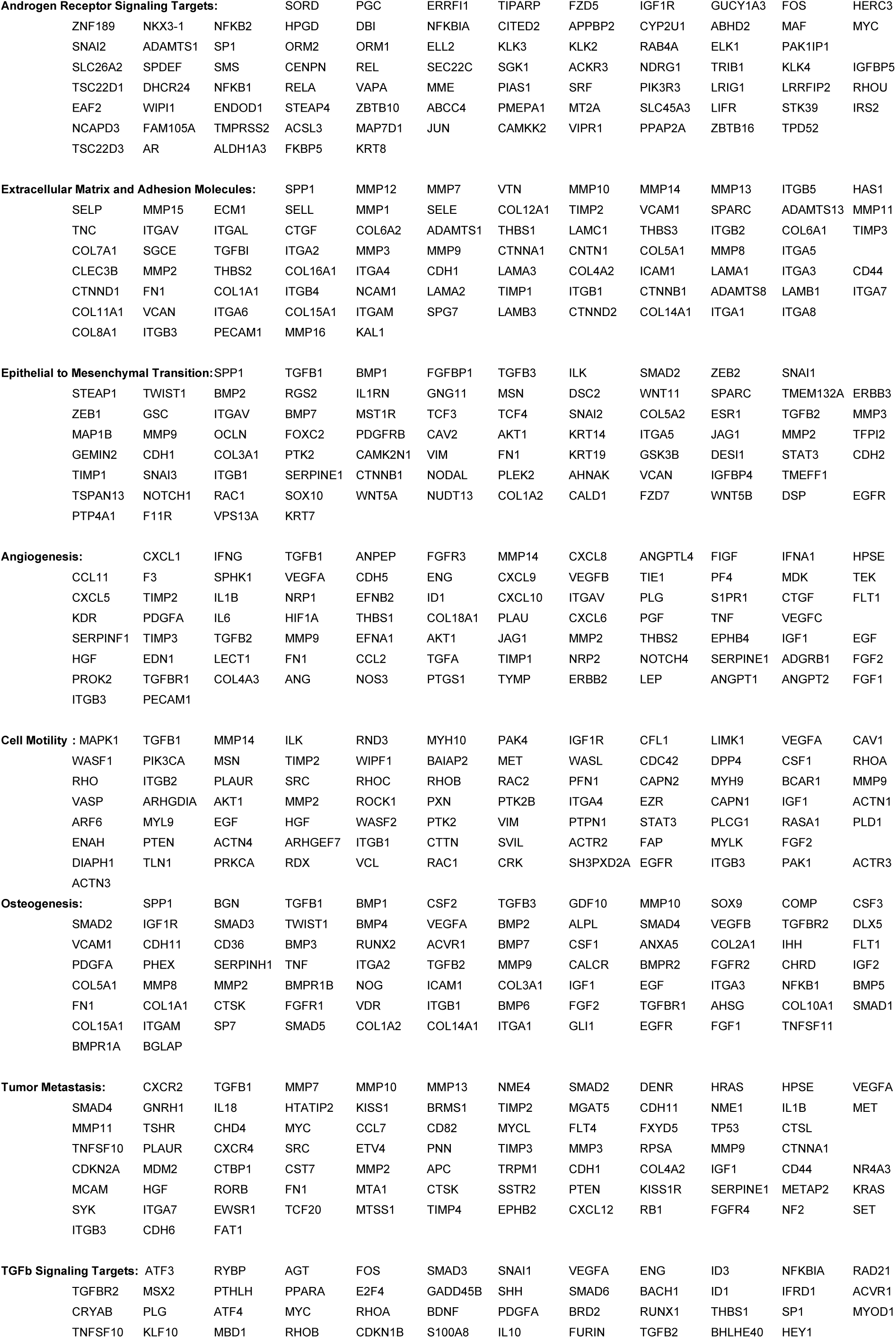
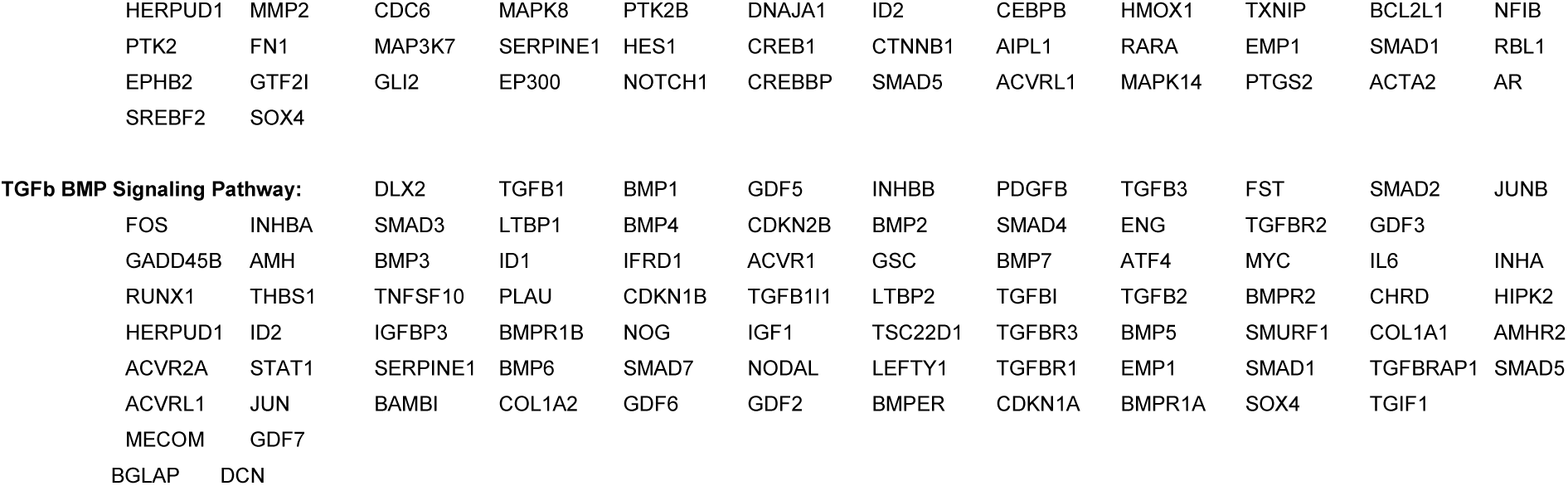
Gene sets used for the correlative analysis.

**Figure S1.**
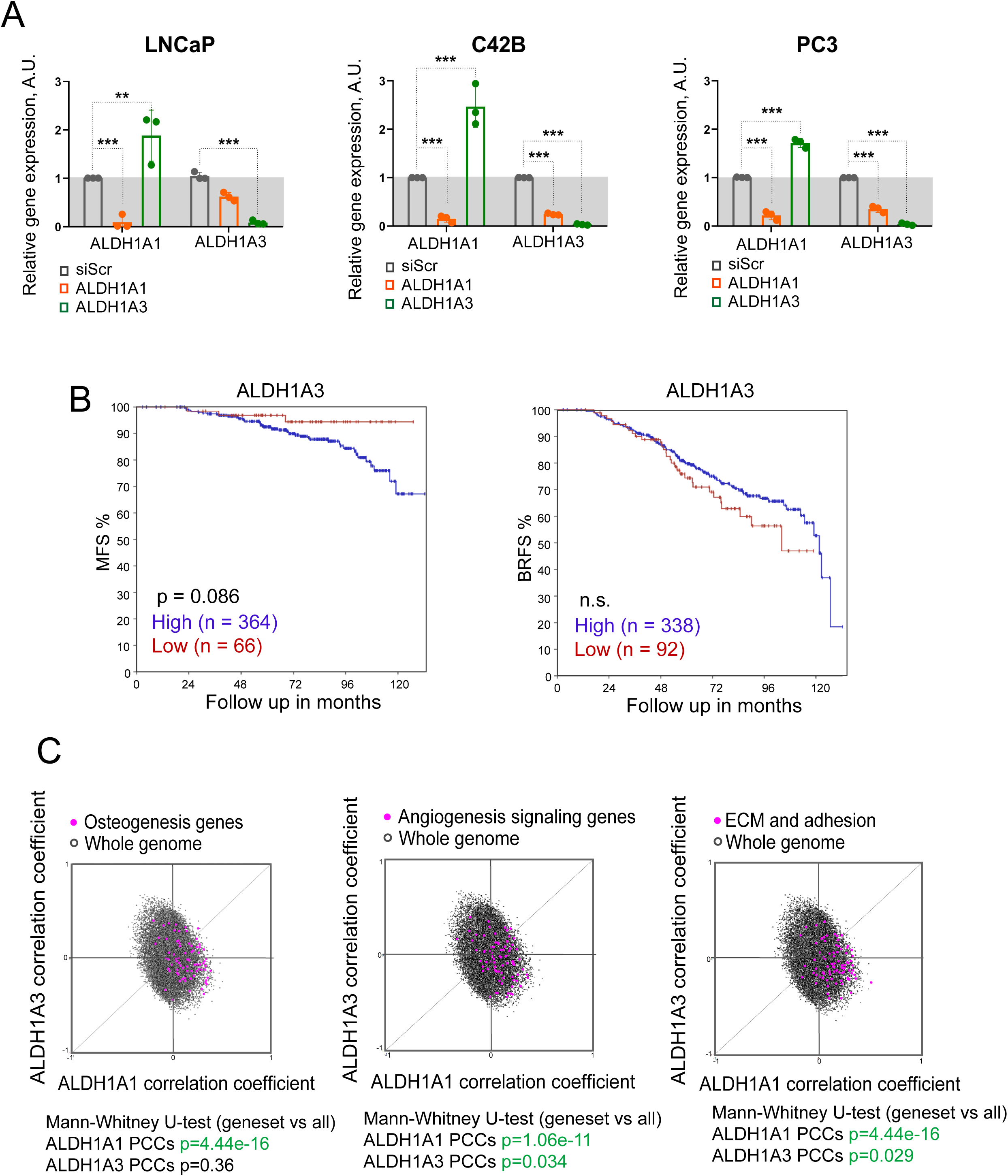
(**A**) Analysis of the ALDH1A1 and ALDH1A3 knockdown. qPCR analysis of the relative ALDH1A1 and ALDH1A3 expression in LNCaP, C42B and PC3 cells upon knockdown of ALDH1A1 or ALDH1A3. Cells transfected with scrambled siRNA (siScr) were used as controls. n = 3; Error bars = SD; **p < 0.01; ***p < 0.001. (**B**) The Kaplan-Meier analyses of the biochemical recurrence-free survival (BRFS) and metastasis-free survival (MFS) for patients with high-risk and locally advanced PCa treated with ADT and then concurrently with one of three radiotherapy regimens as described in Table 3. The patients were stratified by the most significant cut-off for ALDH1A3 expression levels. (**C**) Correlation of mRNA expression levels of ALDH1A1 and ALDH1A3 genes and gene sets related to osteogenesis, angiogenesis, extracellular matrix (ECM) and adhesion in the TCGA PRAD patient cohort (n = 490). The gene lists are provided in Table S2.

**Figure S2.**
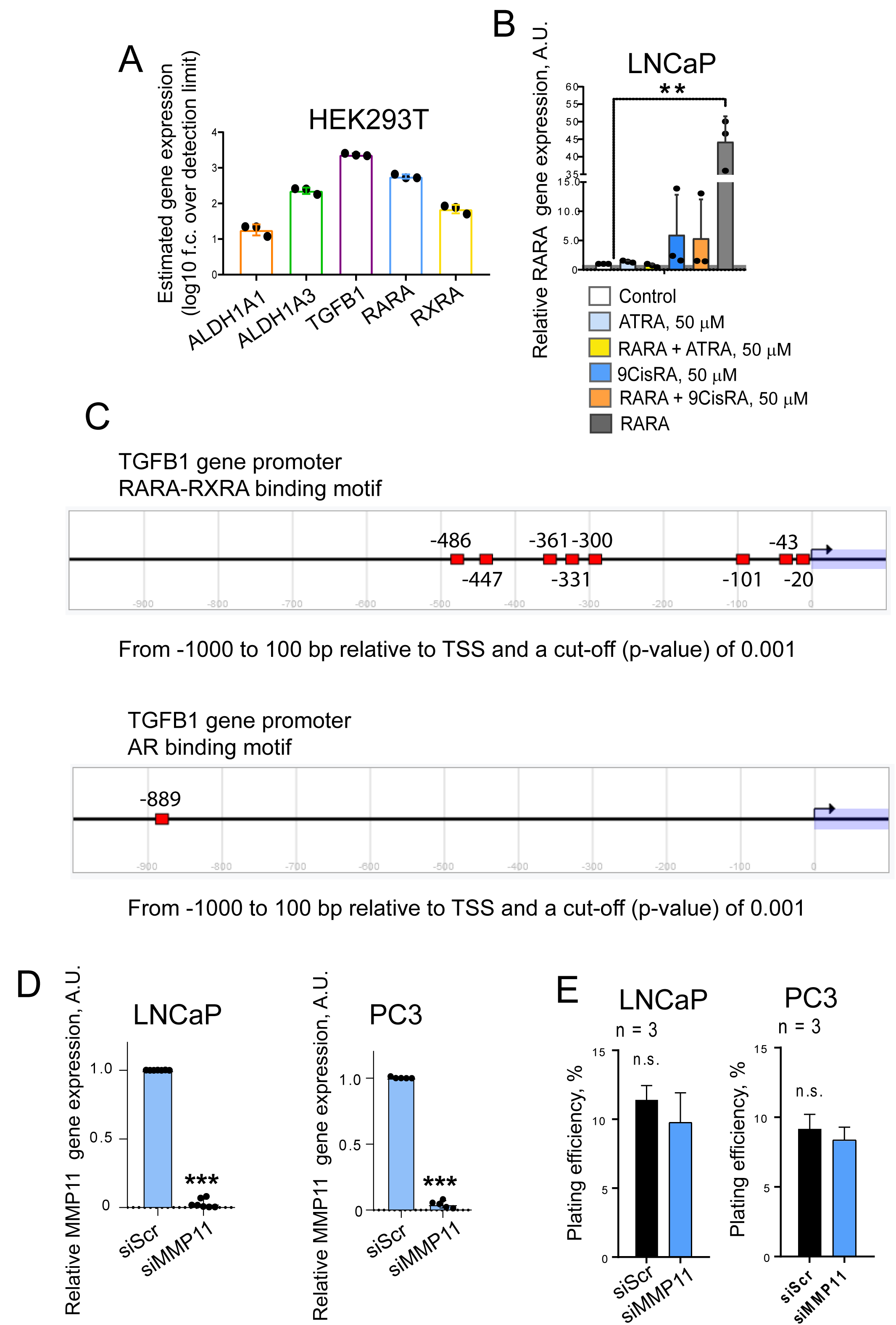
(**A**) qPCR analysis of the relative gene expression in HEK293T cells. n = 3; Error bars = SD. (**B**) qPCR analysis of RARA expression in LNCaP cells upon transient RARA overexpression, treatment with ATRA or 9-cis retinoic acid (9CisRA) for 48 h or both. Cells transfected with empty plasmid were used as control. n = 3; Error bars = SD; **p < 0.01. (**C**) A putative RARA-RXRA and AR binding motifs identified in the TGFB1 gene promoter using the Eukaryotic Promoter Database (https://epd.expasy.org/epd/). (**D**) qPCR analysis of the relative MMP11 expression in LNCaP and PC3 cells upon MMP11. Cells transfected with scrambled siRNA (siScr) were used as controls. n = 3; Error bars = SD; ***p < 0.001. (**E**) Plating efficiencies (PE, %) of LNCaP and PC3 cells after siRNA-mediated knockdown of MMP11. Cells transfected with scrambled (Scr) siRNA were used as controls. n = 3; Error bars = SD. There are no statistically significant differences in plating efficiency among the different conditions (n.s.).

**Figure S3.**
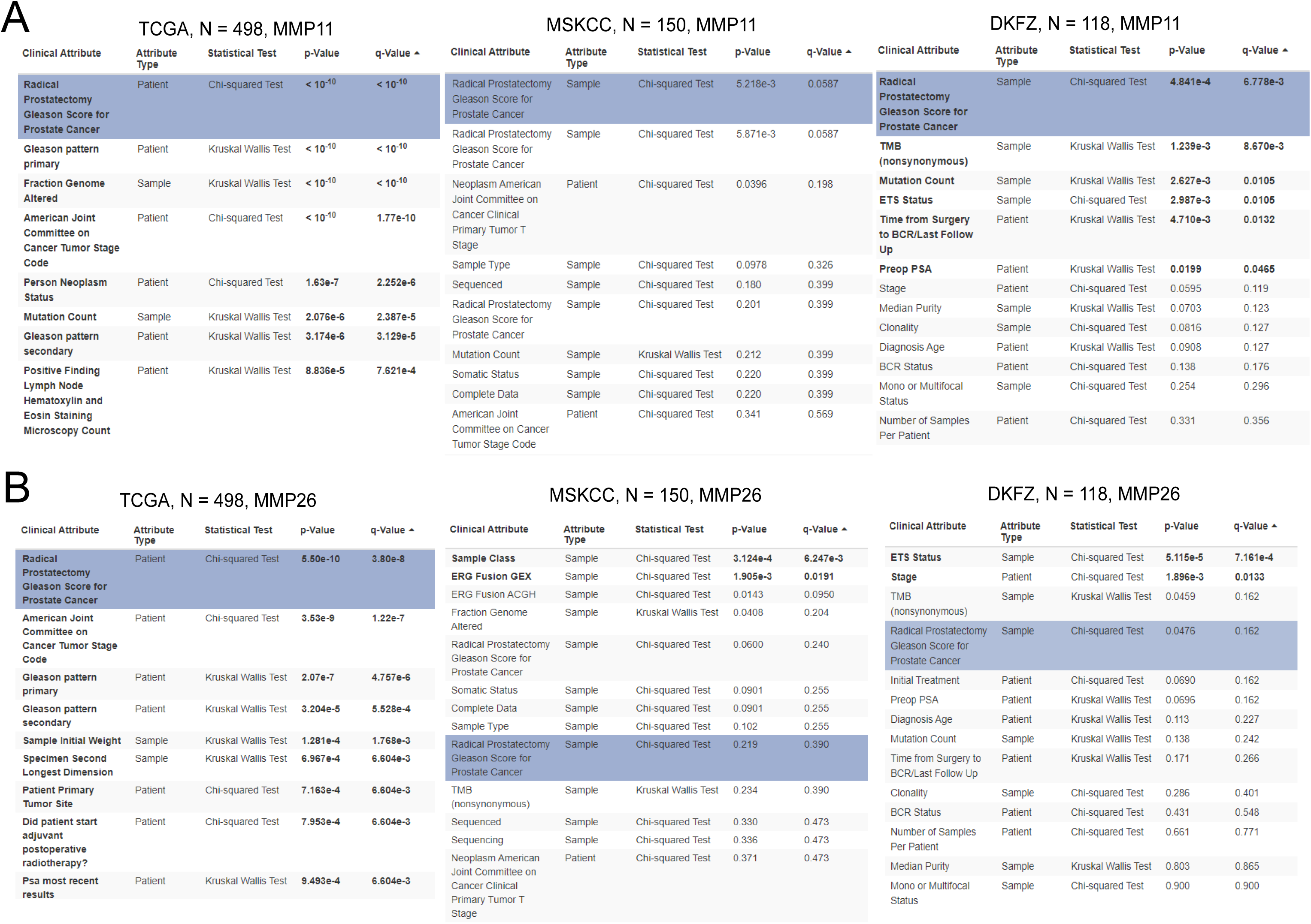
(**A**) Correlation of the MMP11 and (**B**) MMP26 gene expression levels with clinical attributes. Analysis was performed using cBioPortal for cancer genomics (https://www.cbioportal.org).

**Figure S4.**
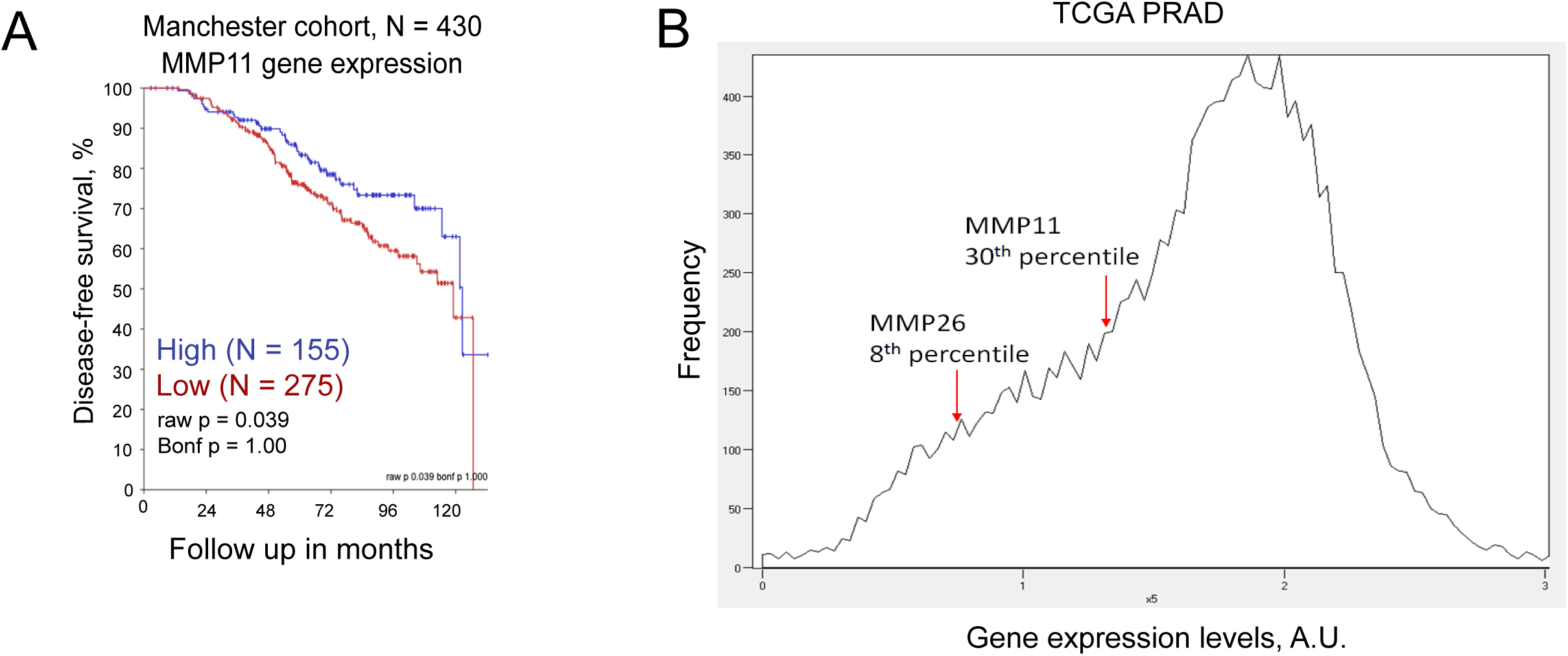
(**A**) The Kaplan-Meier analyses of the association of MMP11 expression and disease-free survival in a retrospective multicenter cohort including patients diagnosed with locally advanced PCa (Manchester dataset, Table 3). (**B**) Distribution of the relative gene expression levels in the TCGA PRAD dataset.

**Figure S5.**
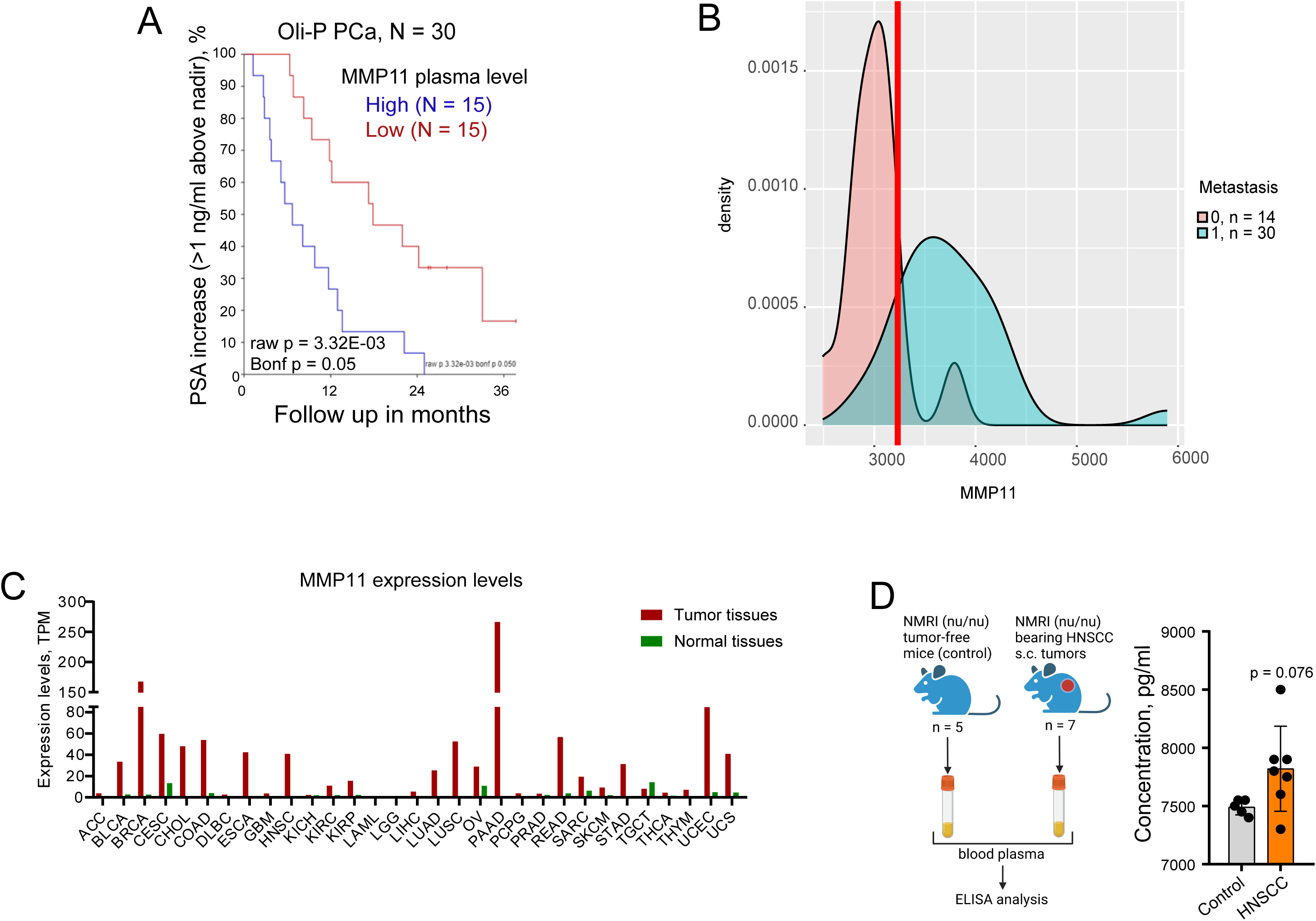
(**A**) The Kaplan-Meier analyses of the association of MMP11 plasma levels and PSA increase (>1 ng/ml above nadir) in patients with oligometastatic PCa treated with local ablative radiotherapy, n = 30. (**B**) A cutoff scan for the ROC analysis is shown in Figure 6D. (**C**) The MMP11 gene expression in tumor samples and normal tissues. The data are obtained using GEPIA 2. TCGA Study Abbreviations: ACC, Adrenocortical carcinoma; BLCA, Bladder Urothelial Carcinoma; LGG, Brain Lower Grade Glioma; BRCA, Breast invasive carcinoma; CESC, Cervical squamous cell carcinoma and endocervical adenocarcinoma; CHOL, Cholangiocarcinoma; COAD, Colon adenocarcinoma; ESCA, Esophageal carcinoma; GBM:Glioblastoma multiforme; HNSC, Head and Neck squamous cell carcinoma; KICH, Kidney Chromophobe; KIRC, Kidney renal clear cell carcinoma; KIRP, Kidney renal papillary cell carcinoma; LAML, acute myeloid leukemia; LIHC, Liver hepatocellular carcinoma; LUAD, Lung adenocarcinoma; LUSC, Lung squamous cell carcinoma; DLBC, Lymphoid Neoplasm Diffuse Large B-cell Lymphoma; OV, Ovarian serous cystadenocarcinoma; PAAD, Pancreatic adenocarcinoma; PCPG, Pheochromocytoma and Paraganglioma; PRAD, Prostate adenocarcinoma; READ, Rectum adenocarcinoma; SARC, Sarcoma; SKCM, Skin Cutaneous Melanoma; STAD, Stomach adenocarcinoma; TGCT, Testicular Germ Cell Tumors; THYM, Thymoma; THCA, Thyroid carcinoma; UCS, uterine carcinosarcoma; UCEC, Uterine Corpus Endometrial Carcinoma; TPM, transcripts per million. (**D**) Analysis of the MMP11 plasma levels in the NMRI (nu/nu) healthy and tumor-bearing mice.

**Figure S6.**
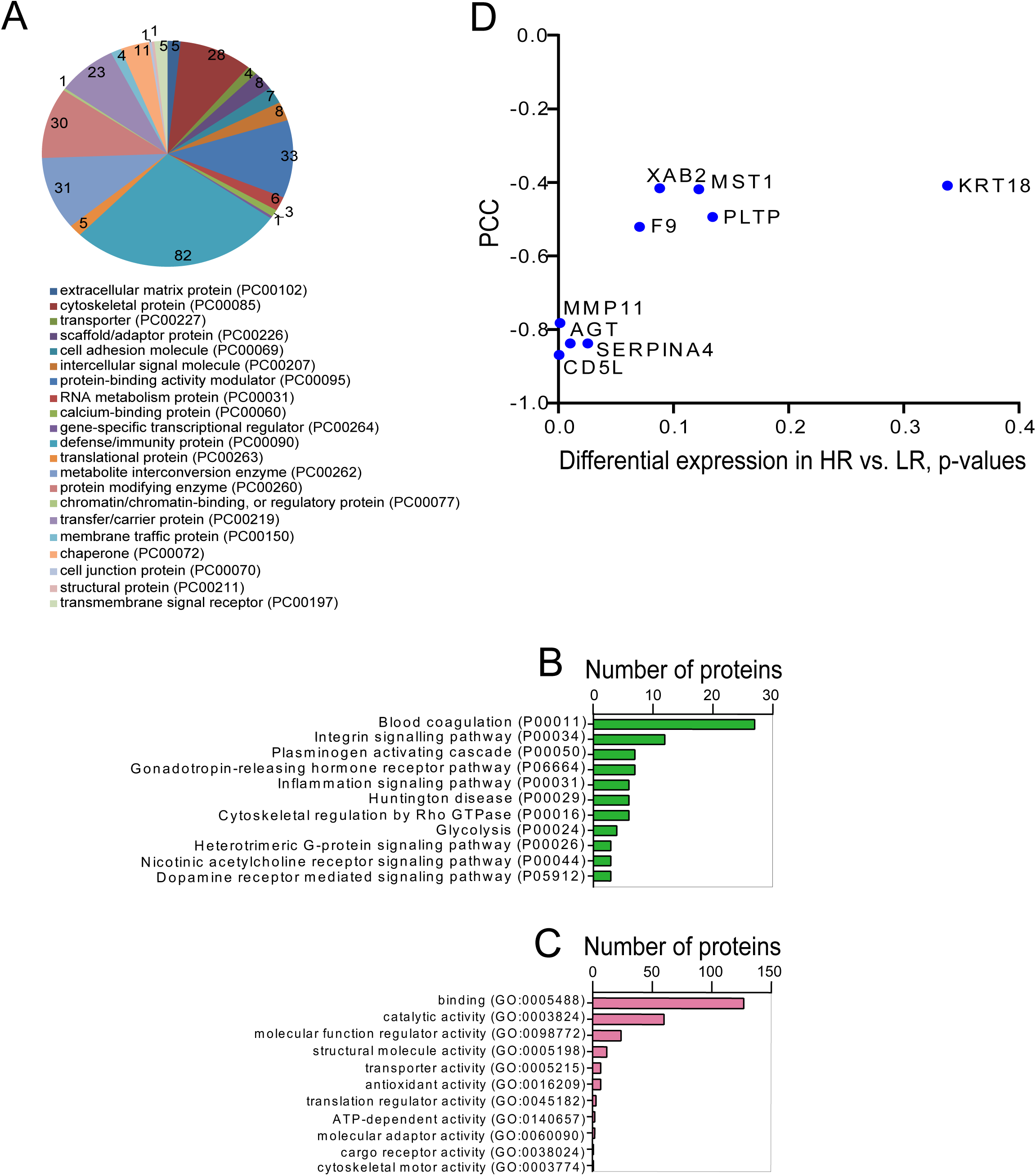
Functional classification of the proteins identified by plasma proteomic profiling (**A**), associated pathways (**B**), and molecular functions according to PANTHER GO-slim (**C**). All analyses were performed using PANTHER18.0 classification system. (**D**) Characterization of 8 selected non-immunoglobulin proteins regarding their comparative expression in a high risk (HR) vs. (LR) groups and correlation with PSA increase (20% above nadir); PCC: Pearson correlation coefficient.

